# B cell receptor silencing reveals the origin of high-grade B cell lymphomas with *MYC* and *BCL2* rearrangements

**DOI:** 10.1101/2024.07.13.603066

**Authors:** Paola Sindaco, Silvia Lonardi, Gabriele Varano, Ilaria Pietrini, Gaia Morello, Piera Balzarini, Hadas Neuman, Filippo Vit, Giorgio Bertolazzi, Hiroshi Arima, Marco Chiarini, Luisa Lorenzi, Vilma Pellegrini, Sabrina Giampaolo, Daniel Garzon, Cecilia Ranise, Mattia Bugatti, Chiara Pagani, Rosa Daffini, Federica Mainoldi, Viveka Selvarasa, Anojan Sivacegaram, Henry Yang, Li Ying, Valeria Cancila, Raoul Bonnal, Euplio Visco, Cristina Lopez Gonzalez, Pasquale Capaccio, Andres J. M. Ferreri, Alessandra Tucci, Antonello D. Cabras, Giancarlo Pruneri, Arianna di Napoli, Reiner Siebert, Riccardo Bomben, Brunangelo Falini, Marco Pizzi, Joo Y. Song, Wing C. Chan, Maurilio Ponzoni, Ramit Mehr, Claudio Tripodo, Fabio Facchetti, Stefano Casola

## Abstract

The B cell receptor (BCR) is essential for mature B cell lymphomas, serving as therapeutic target. Here, we show that high-grade B cell lymphomas with *MYC* and *BCL2* rearrangements (HGBCL-DH-*BCL2*) predominantly exhibit immunoglobulin heavy (IGH) chain silencing, leading to BCR shutdown. HGBCL-DH-*BCL2* with undetectable IGH (IGH^UND^) differ from IGH-expressing counterparts for germinal center-zone gene programs, *MYC* expression and T cell infiltration. While IGH^+^ HGBCL-DH-*BCL2* prefer IGM/IG-Kappa expression, IGH^UND^ counterparts have completed IGH class-switching, favoring IG-Lambda (IGL) light chains. IGH^UND^ HGBCL-DH-*BCL2* preserve *IGHV* gene integrity, overcoming antigen-driven selection. IGH silencing precedes onset and shapes evolution of HGBCL-DH-*BCL2* from Follicular Lymphoma (FL) or FL/HGBCL-DH-*BCL2* common precursor. In FL/HGBCL-DH-*BCL2* pairs and HGBCL-DH-*BCL2* models, BCR silencing promoted RAG1/2-dependent IG light chain editing, causing t(8;22)(q24;q11)/*IGL*::*MYC*. IGH silencing protected HGBCL-DH-*BCL2* models from killing by CD79B-targeting Polatuzumab-Vedotin. Collectively, HGBCL-DH-*BCL2* primarily originate from BCR-silenced isotype-switched t(14;18)/*IGH::BCL2*-positive (pre)FL cells acquiring I*GL::MYC* translocations during IG light chain revision, with clinical implications.

**Significance:** These findings link BCR silencing in isotype-switched t(14;18)^+^ Follicular Lymphoma cells (or their precursors) to RAG1/2 re-expression, promoting *IGL::MYC* translocations responsible for transformation into high-grade B cell lymphomas (HGBCL). Predominant silencing of the BCR complex in HGBCL with *MYC* and *BCL2* rearrangements protects tumor cells from CD79B-directed Polatuzumab-Vedotin killing.

## Introduction

The B cell receptor (BCR) serves as a signaling hub governing the survival and responsiveness of mature B cells to adaptive and innate signals (1). In mature B cell-derived malignancies, the BCR signaling platform is commonly hijacked and otherwise rewired to ensure chronic activation of downstream pathways either autonomously or in response to environmental signals (2,3). Tonic signals from surface (s) BCR also support growth of malignant B cells improving their competitive fitness (4,5). Selection for continuous sBCR expression and signaling in activated B cell-like (ABC) Diffuse large B cell Lymphoma (DLBCL) is achieved via acquisition of somatic mutations targeting the ITAM motif of the CD79B signaling subunit (6,7). Follicular Lymphoma (FL) treated with anti-idiotype antibodies directed against the malignant BCR mutate the target site to escape killing while preserving surface antigen receptor expression (8). In Burkitt lymphoma (BL), the translocation t(8;14)(q24;q32) preferentially selects the non-productive immunoglobulin heavy chain *(IGH)* locus for *IGH*::*MYC* fusions, to preserve BCR expression (9). At the same time, studies in MYC-driven lymphomas have shown that malignant B cells acutely adapt to, and chronically bypass BCR extinction (5,10,11). Similar conclusions were reached through sIG phenotypic investigations in different types of mature B cell neoplasms (12–15). Nonetheless, the concept that mature B cell lymphomas can evolve or eventually arise under conditions of BCR silencing has largely gone unrecognized. The relevance of assessing the influence of BCR silencing on lymphoma genesis and evolution stems from the property of B cells to repeatedly undergo extended periods of antigen receptor downregulation upon participation in the germinal center (GC) reaction. Repeated cell divisions and *IG* variable (*V*) gene hypermutation in the GC dark zone (DZ) area are coupled to extensive BCR turnover, ensuring rapid replacement of antigen receptors carrying germline-encoded IGV genes with mutated counterparts (16). The surface IG^low^ state of GC DZ centroblasts entails the existence of BCR surrogate signals supporting the maintenance of these cells. Whether some B cell lymphomas exploit such mechanisms to gain BCR independence remains unknown. Through a comprehensive phenotypic and molecular screening of more than 300 real-world lymphoma cases with DLBCL morphology, we report recurrent IGH silencing. Lymphomas with undetectable IGH (IGH^UND^) were significantly enriched among GCB-type DLBCL and accounted for the majority of prognosis-poor high-grade B cell lymphomas (HGBCL) with *MYC* and *BCL2* rearrangements (“double-hit”; HGBCL-DH-*BCL2*). Investigations to dissect the mechanisms and consequences of IGH silencing in HGBCL-DH-*BCL2* reveal a mechanistic contribution of antigen receptor silencing to tumor onset and evolution, with potential clinical implications.

## Results

### Dominant IG silencing in high-grade B cell lymphomas with *MYC* and *BCL2* rearrangements

We conducted a screening by immunohistochemistry (IHC) for IGH chains IGM/D/G/A in 253 consecutive B cell non-Hodgkin Lymphomas (B-NHL) cases with DLBCL morphology (Supplementary table S1A). In two thirds of the cases (168/253; 66%), malignant B cells predominantly expressed one IGH chain isotype, primarily IGM, in more than 90% of the cells. In the remaining cases (85/253; 34%), IGH immunoreactivity was undetectable in at least 10% of lymphoma B cells (Fig. 1A-B, Supplementary Fig. S1A, Supplementary table S1A). Among these cases, 56/85 (66%) showed > 90% of atypical B cells with undetectable IGM/D/G/A, hereafter called IGH^UND^ tumors, while the remaining cases (29/85; 34%) consisted of a mosaic of IGH^UND^ (11%-89% of cells) and IGH-expressing tumoral cells, representing the IGH^UND/+^ mixed group (Fig. 1A-B, Supplementary Fig. S1B, Supplementary table S1A). The failure to detect by flow cytometry sIG-kappa (sIGK) or -lambda (sIGL) light chains in representative IGH^UND^ DLBCL cases confirmed and extended IHC results, indicating sBCR silencing (Fig. 1C). IGH^UND^ cases exhibited a significant enrichment for GC B cell-like (GCB) DLBCL as compared to non-GCB counterparts, classified according to the Hans algorithm (Fig. 1D, Supplementary Fig. S1A). Among GCB DLBCL, we observed a preferential association between the IGH^UND^ phenotype and cases subsequently diagnosed as high-grade B cell lymphomas with *MYC* and *BCL2* rearrangements (double-hit, HGBCL-DH-*BCL2*), including a subset carrying also BCL6 rearrangements (HGBCL-DH-*BCL2*-*BCL6*) (Fig. 1E-F). To consolidate these findings, we extended the IGH screening to a total of n=93 HGBCL-DH-*BCL2*(-*BCL6*), scoring 66% with undetectable IGH immunoreactivity, mostly comprising IGH^UND^ cases (Fig. 1G, Supplementary table S1A). The high frequency of IGH^UND^ HGBCL-DH-*BCL2* cases was not solely linked to the common MYC and BCL2 expression in these cases. The HGBCL-DH-*BCL2* cohort (n=93) showed a significantly higher enrichment for IGH^UND^ cases when compared to MYC and BCL2 dual expressor (MB2 DE) GCB DLBCL without rearrangements (n=62) (Fig. 1H, Supplementary table S1B). The striking enrichment for IGH^UND^ cases and the limited understanding of the BCR role in HGBCL-DH-*BCL2*(*-BCL6*) ontogeny and evolution prompted us to investigate the mechanisms and biological implications of IGH silencing in these aggressive lymphomas.

**Figure 1.**
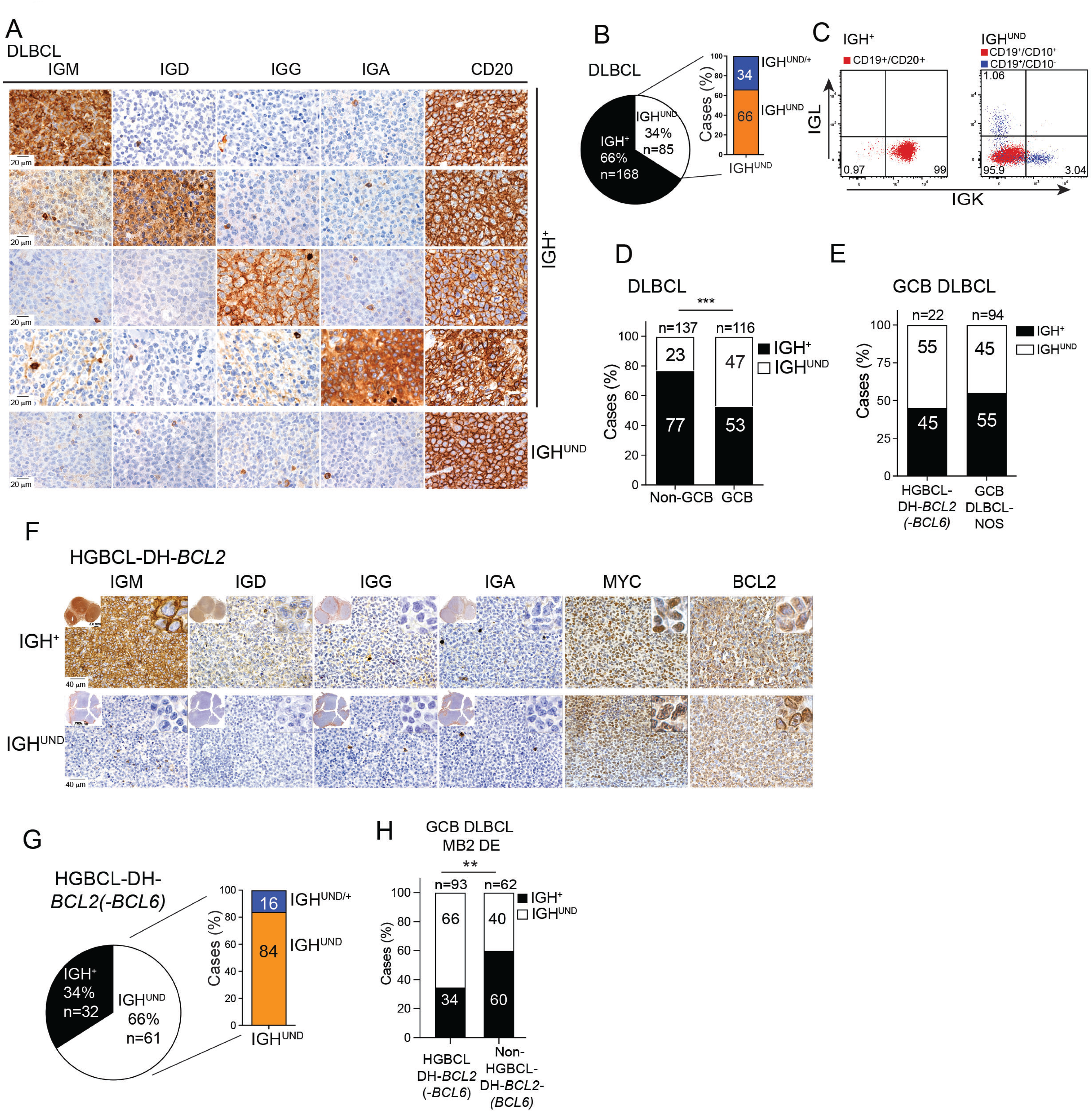
IGH silencing in GCB DLBCL and HGBCL-DH-*BCL2*. (A) Class-specific IGH IHC analysis of representative DLBCL cases, including an IGH^UND^ case (last row). CD20 expression and scattered IGH immunoreactivity in surrounding plasma cells acted as internal staining controls. (B) Distribution of IGH^+^ and IGH^UND^ lymphomas among consecutive DLBCL cases (n=253). The IGH^UND^ group includes IGH^UND/+^ mixed lymphomas. Numbers inside bars indicate corresponding frequencies. (C) FACS analysis for sIGK and sIGL expression in lymph node cell suspensions of representative IGH^+^ and IGH^UND^ DLBCL. Percentage of gated tumoral (red) and normal B cells (blue) are shown. (D) Frequencies of IGH^+^ and IGH^UND^ cases among DLBCL cases according to the Hans algorithm (***p<0.001; Fisher’s exact test). (E) Frequencies of IGH^+^ and IGH^UND^ cases among HGBCL-DH-*BCL2(-BCL6)* and GCB DLBCL NOS cases. Number of cases analyzed is indicated above histograms. (F) Class-specific IGH IHC in representative IGH^+^ and IGH^UND^ HGBCL-DH-*BCL2* cases with infiltrating plasma cells acting as internal staining control. Insets represent sections photographed at low (upper left,) and high (upper right) magnification, respectively. (G) Frequencies of IGH^+^ and IGH^UND^ cases among HGBCL2-DH-*BCL2(-BCL6)* of an extended cohort of n=93 cases. The IGH^UND^ group included a minor fraction of IGH^UND/+^ mixed lymphomas. (H) Frequencies of IGH^+^ and IGH^UND^ cases among GCB MB2 DE DLBCL, clustered according to the presence (HGBCL2-DH-*BCL2*) or absence (non-HGBCL-DH-*BCL2*) of *BCL2* and *MYC* rearrangements (**p<0.01; Fisher’s exact test).

### IGH^UND^ HGBCL2-DH-*BCL2* boost *MYC* expression and limit T cell engagement

We conducted a bulk transcriptomics comparison of IGH^+^ and IGH^UND^ HGBCL-DH-*BCL2*(-*BCL6*) cases (n=14), including in the analyses also a subset of MB2 DE GCB DLBCL cases (n=8; Supplementary table S2A-B). Unsupervised clustering based on the full transcriptome profile mostly separated IGH^UND^ lymphomas from IGH^+^ counterparts (Fig. 2A) suggestive of distinct biological entities. IGH^UND/+^ HGBCL-DH-*BCL2*(*-BCL6*) fell within the IGH^UND^ cluster and were thus incorporated into the latter group for downstream analyses. IGH^UND^ tumors differed from their IGH^+^ counterparts for an almost equal number of up- (n=468) and down-regulated (n=485) genes (log2FC > 0.58; p-adj value <0.05) (Fig. 2B and Supplementary table S2C). IGH^UND^ tumors showed significantly (p-adj value <0.05) higher expression of *MYC* (Fig. 2C), cell-cycle regulators, and factors involved in DNA replication and repair, mitosis, chromatin remodeling, telomere maintenance, mitochondrial respiration, and sugar, nucleotide, and amino acid metabolism (Fig. 2D, Supplementary table S2D). IGH^UND^ lymphomas also expressed higher transcripts for components of the unfolded protein stress response and the SKP1-cullin 1(CUL1)-F-box E3 ligase complex (Fig. 2D and Supplementary table S2D), suggesting a tight control over protein turnover. IGH^+^ lymphomas showed stronger expression of interferon gamma-, IRF8- and NF-κB- regulated genes and immunomodulatory factors such as IL10, TGFB3, IDO1, IDO2 and CCL22 (Fig. 2D and Supplementary table S2D). To interrogate the HGBCL-DH-*BCL2* tumor ecosystem at higher resolution, distinct regions of interest (ROI) of representative IGH^+^ and IGH^UND^ cases were profiled by spatial transcriptomics for 1825 cancer- and immune-relevant genes (Fig. 2E, Supplementary Fig. S2A and Supplementary table S2E). IGH^UND^ HGBCL-DH-*BCL2* ROI showed significant up- (n=234) and down-(n=260) modulation of transcripts compared to IGH^+^ counterparts (Fig. 2E and Supplementary table S2E; p-adj value <0.05). In IGH^+^ HGBCL-DH-*BCL2* ROI, higher expression of several chemokines, adhesion molecules and extracellular matrix proteins suggested a richer representation of immune, inflammatory, and stromal/vascular cell types (Fig. 2E, Supplementary table S2F). IGH^+^ HGBCL-DH-*BCL2* tumor niches showed higher expression of genes associated with follicular dendritic cells, NK cells, and myeloid cells (Supplementary table S2F), in line with cell-type deconvolution analyses (Supplementary Fig. S2B). Transcripts for T cell receptor complex components, T-Follicular Helper, T_H_17 cells and T-regulatory cells were also more represented in IGH^+^ HGBCL-DH-*BCL2* ROI (Supplementary table S2F). IHC analyses confirmed a richer infiltration of CD3^+^ T cells in IGH^+^ HGBCL-DH-*BCL2*(*-BCL6*), as compared to the IGH^UND^ counterparts (Fig. 2F and Supplementary table S2G). Conversely, IGH^UND^ HGBCL-DH-*BCL2* ROI showed stronger expression of *MYC,* and factors involved in redox regulation, mitochondrial respiration, cell cycle progression, DNA replication and repair (Fig. 2E; Supplementary table S2F), in agreement with bulk transcriptomics data. Collectively, IGH^UND^ and IGH^+^ HGBCL-DH-*BCL2* establish distinct tumor microenvironments, with strongest *MYC* expression and preferential T cell evasion in IGH-silenced cases.

**Figure 2.**
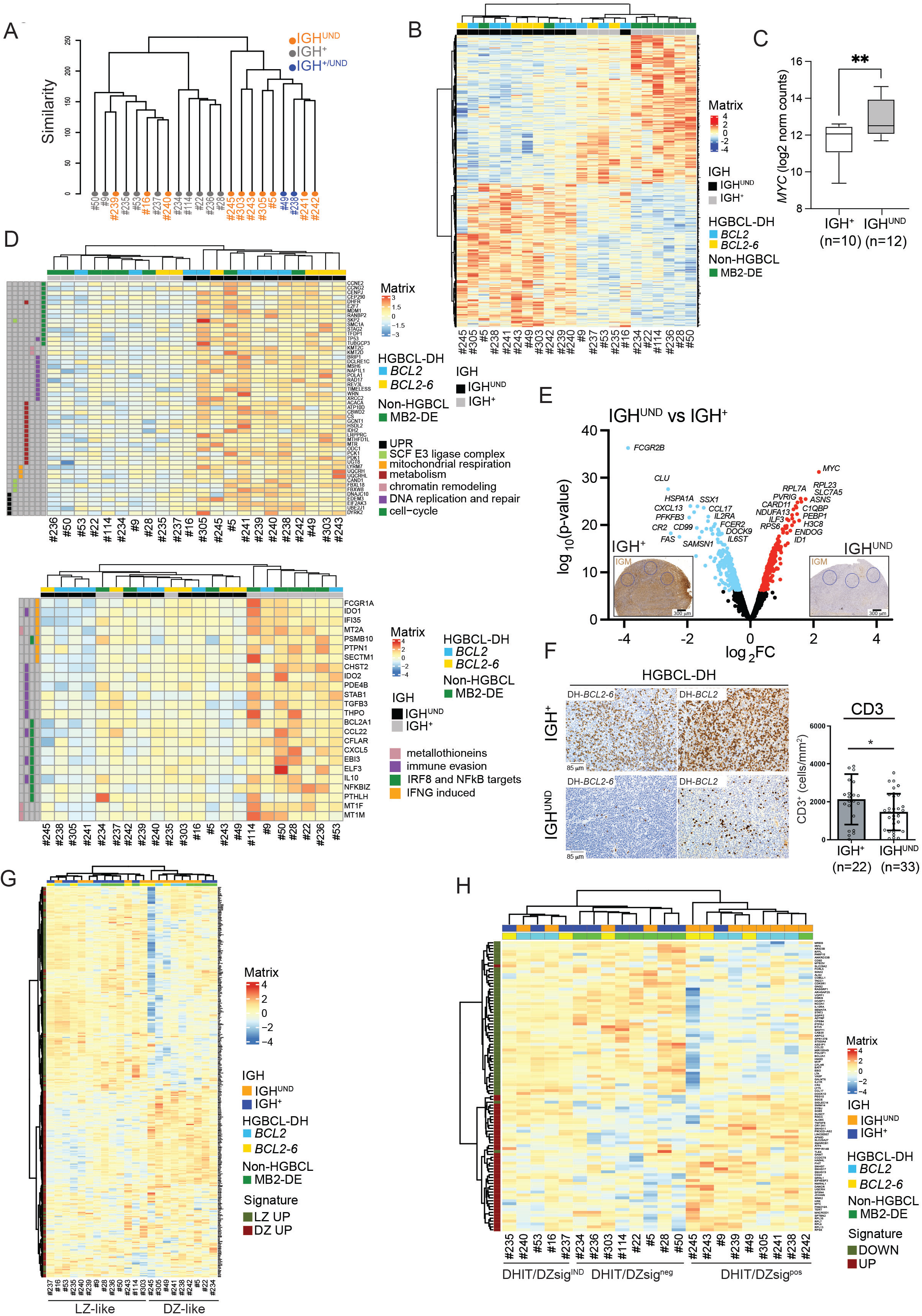
IGH^+^ and IGH^UND^ HGBCL-DH-*BCL2* differ in gene expression and T cell infiltrate. (A) Unsupervised clustering of GCB MB2 DE DLBCL cases (n=22) based on whole transcriptome data. Case ID is listed below. Cases were clustered for IGH expression (grey) or IGH^UND^ (red) state (red) IGH^UND/+^ mixed cases are shown in blue. (B) Heatmap representation of differentially expressed genes (n=953; log2 FC>0.58: adj p-value<0.05) between IGH^+^ and IGH^UND^ MB2 DE DLBCL, after unsupervised clustering. IGH and HGBCL-DH-*BCL2* status is indicated above the heatmap. Z-score normalized expression values are shown. (C) Boxplot of median *MYC* transcripts (horizontal line) and 5th-95th percentile (whiskers) in GCB MB2 DLBCL (*n*=22) clustered according to IGH expression (**p<0.01; unpaired t-test). (D) Unsupervised clustering of MB2 DLBCL cases for gene transcripts respectively up- (upper map) and down- (lower map) regulated in IGH^UND^ compared to IGH^+^ subset, grouped based on gene ontology. (E) Volcano plot representation of differentially expressed genes (log2 FC >0.58: adj p-value <0.05) between ROI of IGH^UND^ and IGH^+^ HGBCL-DH-*BCL2*. Top differentially expressed genes are shown. (F) CD3 IHC analysis in representative IGH^+^ and IGH^UND^ HGBCL-DH-*BCL2*(-*BCL6*) cases. Histograms refer to mean number of CD3^+^ cells/mm^2^ +/- standard deviation (p<0.05; unpaired t-test with Welch’s correction). Circles represent individual cases. (G) Unsupervised clustering of IGH^+^ and IGH^UND^ MB2 DLBCL cases for GC spatial DZ/LZ signatures. Heatmap identifies genes preferentially expressed in DZ (red) or LZ (green) regions. For each case, IGH and HGBCL-DH-*BCL2(-BCL6)* status are indicated above the heatmap. (H) Unsupervised clustering of IGH^+^ and IGH^UND^ MB2 DE DLBCL cases according to the DHIT/DZ molecular signature. Gene transcripts positively (red) and negatively (green) associated with the DHIT/DZ signature are indicated.

### IGH^UND^ and IGH^+^ HGBCL-DH-*BCL2* resemble different germinal center B cell subsets

We recently established GC DZ and LZ spatial gene expression signatures (Supplementary table S2H) of polarized reactive GCs from human tonsil (Supplementary table S2H, (25)). Unsupervised interrogation of RNA-seq data for GC spatial signatures identified two main clusters of GCB-type MB2 DE lymphomas (Fig. 2G). The first (n=9; 41%), mostly comprising IGH^UND^ cases (7/9, 78%), more closely resembled GC DZ B cells (hereafter called “DZ-like”). The second cluster (n=13; 59%), consisting primarily of IGH^+^ lymphomas (8/13; 61%), showed closer transcriptional relationship to GC LZ B cells (“LZ-like”; Fig. 2G). DZ-like HGBCL-DH-*BCL2*(*-BCL6*) were entirely included within the IGH^UND^ subset, while LZ-like HGBCL-DH-*BCL2*(*-BCL6*) included both IGH^+^ tumors and IGH^UND^ cases. Assignment of HGBCL-DH-*BCL2*(*-BCL6*) cases to DZ- or LZ-like clusters was confirmed interrogating independent GC DZ- and LZ B cell signatures (Supplementary Fig. S2C, (23)). Spatial transcriptomics of HGBCL-DH-*BCL2* indicated preferential expression of DZ-associated transcripts in ROI of the IGH^UND^ subset, while IGH^+^ counterparts expressed preferentially LZ-localized mRNAs (Supplementary Fig. S2D). Interrogation of bulk transcriptome data for the Double-Hit/DZ molecular signature (DHIT/Dzsig (24,53)) identified a DHIT/DZsig^+^ group mostly consisting of IGH^UND^ HGBCL-DH-*BCL2*(*-BCL6*) (8/9; 89%). The remaining cases, including both IGH^+^ (n=3) and IGH^UND^ (n=2) HGBCL-DH-*BCL2*(*-BCL6*) showed a DHIT/DZ “indeterminate” signature with concurrent expression of DHITsig up- and down-regulated genes (Fig. 2H). In line with previous evidence (24), DHIT/DZsig-negative MB2 DE lymphomas mostly lacked *BCL2* rearrangements (6/8 cases; 75%) while conserving IGH expression (7/8 cases; 87%) (Fig. 2H).

### Shared and private mutational landscape of IGH^UND^ and IGH^+^ HGBCL-DH-*BCL2*

We compared the mutational landscape of IGH^UND^ and IGH^+^ HGBCL-DH-*BCL2*(*-BCL6*), performing whole exome sequencing (Supplementary table S3). The analysis was limited to non-synonymous single nucleotide variants (nsSNV) of genes reported to be mutated in DLBCL, FL, HGBCL-DH-*BCL2*, and BL, to counterbalance the lack of matching germline DNA (Fig. 3A and Supplementary table S3A). In both IGH^+^ and IGH^UND^ HGBCL-DH-*BCL2*(*-BCL6*) cases, nsSNV were found in genes coding for chromatin remodelers (*CREBBP*, *KMT2C/D*, *EP300/400*), linker Histone *H1 D-E* variants, and *BCL7A*, suggestive of a common origin from classical FL (Fig. 3A and Supplementary table S3). Non-silent mutations in genes relevant for GC B cell biology (*TNFRSF14, GNA13, KLHL14, SOCS1*) were also identified in both IGH^+^ and IGH^UND^ HGBCL-DH-*BCL2*(*-BCL6*), along with human leukocyte antigen (HLA) class-1 inactivating mutations (*HLA-A, HLA-B, HLA-C*) (Fig. 3A and Supplementary table S3). Irrespective of IGH status, HGBCL-DH-*BCL2*(*-BCL6*) carried nsSNV in apoptosis and cell-cycle genes, including *TP53, BCL2, BIRC6, CCND3 and CDKN2A* (Fig. 3A and Supplementary table S3). Identification of nsSNVs in DNA damage checkpoint kinases *ATM and ATR* and genes involved in DNA mismatch (*MSH2, MSH4, MSH6, MSH5, MLH1*), nucleotide excision (*ERCC2, ERCC3, ERCC4, ERCC5, ERCC6*), and homologous recombination (*BRCA1, BRCA2*) repair, and in DNA polymerases (*POLB, POLE/E2*) suggest a general status of genome instability and DNA hypermutation in both IGH^+^ and IGH^UND^ tumor subsets (Fig. 3A and Supplementary table S3). On such shared mutational background, IGH^UND^ and IGH^+^ HGBCL-DH-*BCL2*(*-BCL6*) selected distinct nsSNVs. IGH^UND^ lymphomas exhibited a preference for mutations in genes regulating MYC protein levels (i.e., *MYC^T58A^*; *FBXW7*(*54*)) (Fig. 3A and Supplementary table S3). Gain-of-function mutations in *EZH2* (Y646N, A692V) and *FOXO1* (M1L, R21H, T24I, C23W) were identified in IGH^UND^ HGBCL-DH-*BCL2*(*-BCL6*), along with mutations in genes regulating the GC DZ B cell program (*BCL6, MEF2B, IRF8, KLHL6*) (Fig. 3A and Supplementary table S3). N-terminus *SGK1* truncating mutations and a *BRAF^G469A^* gain-of-function (55) mutation suggested respectively heightened PI3K signaling (56) and constitutive RAS/MAPK activation specifically in IGH^UND^ HGBCL-DH-*BCL2*(*-BCL6*) cases (Fig. 3A and Supplementary table S3). While mutations in JAK/STAT signaling genes *SOCS1* and *TYK2* were shared between IGH^+^ and IGH^UND^ HGBCL-DH-*BCL2*(*-BCL6*), *STAT6* mutations were unique to IGH^+^ tumors (Fig. 3A and Supplementary table S3). Bi-allelic *TET2* nsSNVs were identified in one IGH^+^ HGBCL-DH-*BCL2* case, suggesting interference with DNA methylation. Also, mutations in effectors of the TLR4/NF-κB signaling pathway (*TLR4, IRAK2, IRAK3, RIPK3, NFKBIB*) were uniquely observed across IGH^+^ tumors (Fig. 3A and Supplementary table S3). These results indicate that IGH^+^ and IGH^UND^ HGBCL-DH-*BCL2*(*-BCL6*) share a set of genetic alterations possibly inherited from a preceding FL (or common precursor), from which they diverge following distinct evolutionary trajectories.

**Figure 3.**
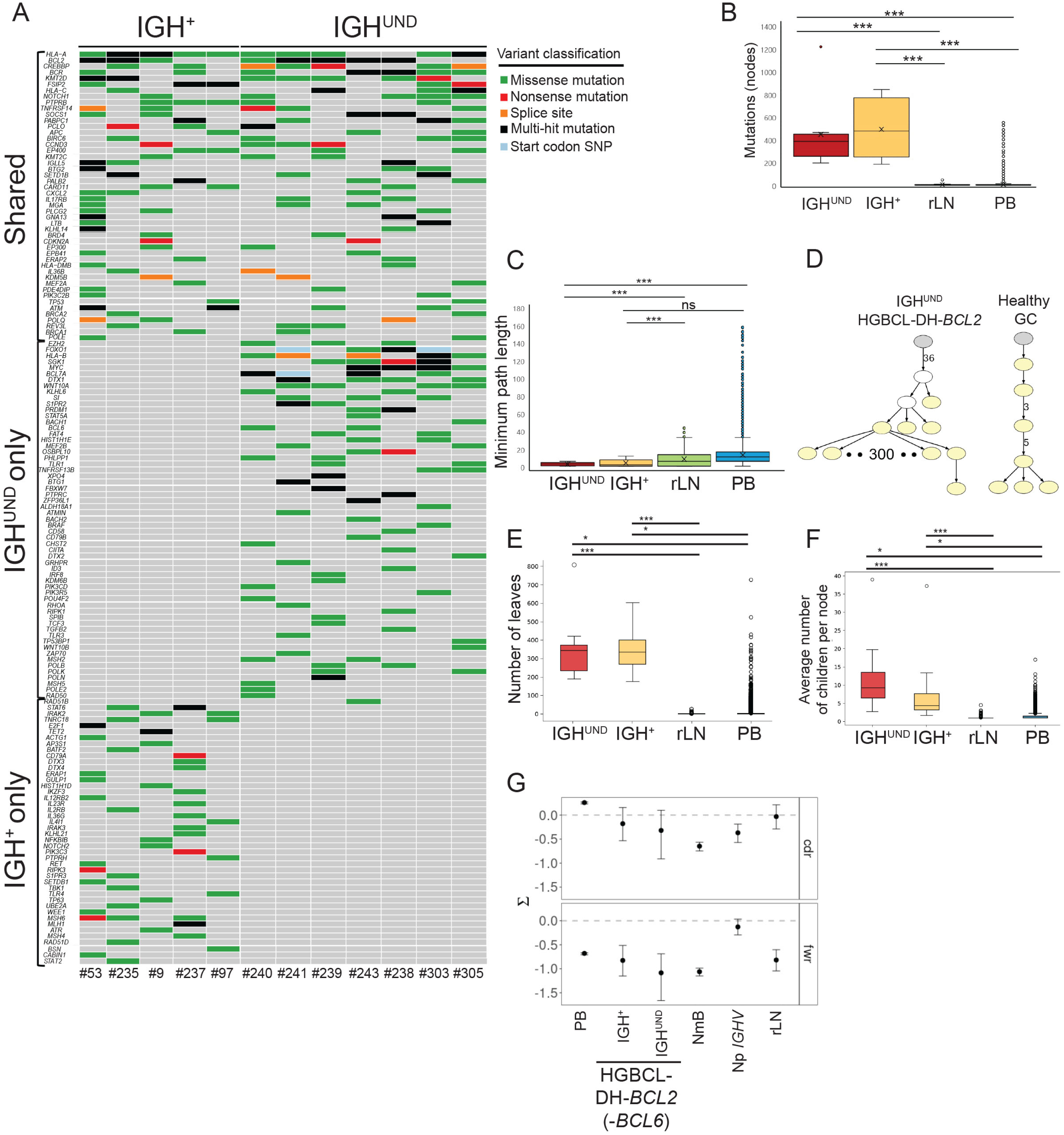
Mutational analyses in IGH^+^ and IGH^UND^ HGBCL-DH-*BCL2*(-*BCL6*) (A) Nonsynonymous single nucleotide variants for selected genes in IGH^+^ and IGH^UND^ HGBCL-DH-*BCL2*(-*BCL6*), or both. Colors indicate gene variant features. (B) Total number of mutations in *IGHV* rearrangements of HGBCL-DH-*BCL2* (-*BCL6*), peripheral blood (PB) and reactive lymph node (rLN) healthy B cell clones, counted by IgTreeZ-MTree. ***Mann Whitney U test with Benjamini-Hochberg correction for multiple comparisons, p-value<0.001. (C) Minimum root to leaf path corresponding to the minimum number of mutations per sequence in *IGHV* rearrangements of HGBCL-DH-*BCL2*, PB and rLN B cell clones, measured by IgTreeZ-MTree. ***Mann Whitney U test with Benjamini-Hochberg correction for multiple comparisons, p-value<0.001. (D) Representative lineage tree of an IGH^UND^ HGBCL-DH-*BCL2* clone. Yellow filled nodes represent sampled sequences. Numbers on edges indicate numbers of mutations between nodes; edges without a number represent one mutation. A lineage tree from a healthy GC B cell clone is also shown for comparison. (E) Numbers of leaves per dominant clone for the indicated samples, counted by IgTreeZ- MTree. A * or ***, Mann Whitney U test with Benjamini-Hochberg correction for multiple comparisons, p-value<0.05 or <0.001. (F) Average number of children per node for the indicated samples counted by IgTreeZ-MTree. Mann Whitney U test with Benjamini-Hochberg correction for multiple comparisons, p-value<0.05 (*) or <0.001 (***). (G) Mean and confidence intervals of selection scores (Σ) for CDR and FRW regions of clonal *IGHV* rearrangements from HGBCL-DH-*BCL2(-BCL6)* cases, antigen-selected PB B cells and rLN B cells from healthy donors, as calculated by BASELINe. Selection scores were also calculated for clonal rearrangements of HGBCL-DH-*BCL2*(*-BCL6*)-associated non malignant B cells (NmB). Non productive (Np) IGHV rearrangements from healthy GC-experienced B cell clones were included in the analyses as non-selected controls.

### IGH^UND^ HGBCL-DH-*BCL2* preserve productive *IGHV* rearrangements

High-throughput sequencing of *IGHV* gene libraries identified single-clone rearrangements in 22 HGBCL-DH-*BCL2*(*-BCL6*) cases (Supplementary table S3B). Together with IGH^+^ tumors, all IGH^UND^ HGBCL-DH-*BCL2*(*-BCL6*) cases carried potentially productive, hypermutated clonal *IGHV* rearrangements (Supplementary table S3B). We compared the mutational pattern of *IGHV* rearrangements from IGH^UND^ HGBCL-DH-*BCL2*(*-BCL6*) to that of GC-experienced B cell clones from reactive lymph nodes (rLN) and peripheral blood (PB) of healthy individuals, as reference. For tumor *IGHV* analyses, we focused on mutations acquired after B cell transformation, excluding mutations on lineage tree trunks. For healthy B cell clones, we included trunk mutations to capture antigen-driven selection features, reaching similar results restricting analyses to trunk-less counterparts (data not shown). Malignant GC B cells in IGH^UND^ HGBCL-DH-*BCL2*(*-BCL6*) exhibited significantly more mutations per clone than healthy counterparts (Fig. 3B), compatible with ongoing AID expression detected in most cases (Supplementary Fig. S3A and Supplementary table S3C). The minimal numbers of mutations per individual *IGHV* sequence, however, was lower in IGH^UND^ lymphomas compared to healthy GC/post-GC B cells, excluding expansion of particular subclonal variants (Fig. 3C). This was consistent with lineage tree shapes of IGH^UND^ HGBCL-DH-*BCL2* clones, indicating weaker vertical growth compared to healthy counterparts (Fig. 3D). Lineage tree topology analysis suggested loosened antigen-driven selection in IGH^UND^ HGBCL-DH-*BCL2*(*-BCL6*), reflected in significantly more leaves (i.e., cells without descendants) and higher average number of children per node compared to rLN B cells (Fig. 3E-F). We applied the BASELINe algorithm (45) to estimate *IGHV*-driven selection strength in HGBCL-DH-*BCL2*(*-BCL6*), separately analyzing CDR and FWR for patterns of mutations. IGH^UND^ HGBCL-DH-*BCL2*(*-BCL6*) preserved selection against amino acid replacement (R) mutations within FWRs (Fig. 3G), with scores for CDRs remaining neutral to weakly negative. *IGHV* rearrangements from circulating healthy IG-mutated antigen-selected B cell clones in healthy controls showed the expected selection for R mutations in CDR and against R mutations in FWR. Such selection was missing in clones bearing out-of-frame rearrangements, confirming unbiased accumulation of R mutations. Negative selection for R mutations in CDRs of GC B cell clones from rLN and from lymphoma-associated non-malignant B cells suggested achievement of optimal BCR binding to cognate antigen, counter selecting further amino acid substitutions. Altogether, IGH^UND^ HGBCL-DH-*BCL2*(*-BCL6*) preserve functional integrity of, and refrain from structurally disrupting IGHV domains, while overcoming antigen-driven selection.

### IGH^UND^ and IGH^+^ HGBCL-DH-*BCL2* differ in IGH class choice

We tested whether IGH shutdown was linked to a particular IGH class selected by the HGBCL-DH-*BCL2*(*-BCL6*) precursor B cell. Analysis of RNA-seq data of IGH^+^ HGBCL-DH-*BCL2*(*-BCL6*) cases indicated predominant expression of *IGHM* constant region mRNAs, concordant with IHC data (Fig. 4A, Supplementary table S4A). In IGH^UND^ HGBCL-DH-*BCL2*(*-BCL6*), low/undetectable *IGHM* was replaced by class switched *IGHG1* (n=2), *IGHG2* (n=2), *IGHG4* (n=2), *IGHE* (n=1) or *IGHA_1/2_* (n=3) transcripts (Fig. 4A, Supplementary table S4A). Notably, compared to IGH^+^ HGBCL-DH-*BCL2*(*-BCL6*), *IGH* mRNA levels were significantly reduced in the IGH^UND^ subset, pointing to reduced steady-state transcripts contributing to IGH silencing (Fig. 4B). To exclude interference from tumor-infiltrating plasma cells expressing high *IGH* mRNA levels, we tracked in situ *IGHM/D/G/A* transcripts by RNA scope technology (n=30). In accordance with RNA-seq data, 10/11 IGH^+^ HGBCL-DH-*BCL2*(*-BCL6*) B cells expressed *IGHM* (occasionally with *IGHD)* transcripts. In contrast, IGH^UND^ HGBCL-DH-*BCL2*(*-BCL6*) malignant B cells expressed either *IGHG (*n=17) or *IGHA (*n=2) mRNAs, at the expense of *IGHM/D* transcripts, which were largely undetectable (Fig. 4C and Supplementary table S4B). To establish whether HGBCL-DH-*BCL2*(*-BCL6*) experienced IGH class switch recombination (CSR), quantitative genomic PCR for a segment of the *IGHM* gene, which is lost after IG CSR, was carried out in representative cases. Differently from healthy naive IGM^+^ B cells carrying two *IGHM* loci, in 7/8 (88%) IGH^UND^ HGBCL-DH-*BCL2*(*-BCL6*), *IGHM* gene copy number (GCN) dropped below the unit, consistent with bi-allelic IG CSR, with residual *IGHM* alleles likely contributed by tumor-infiltrating healthy IGM^+^ B/plasma cells (Fig. 4D, Supplementary table S4C). All IGM^+^ HGBCL-DH-*BCL2*(*-BCL6*) carried on average one *IGHM* GCN (Fig. 4D, Supplementary table S4C), compatible with monoallelic IG CSR on the non-productive *IGH* chromosome(57). In summary, whereas IGH^UND^ HGBCL-DH-*BCL2*(*-BCL6*) originate from IGG/A-switched B cells, most IGH^+^ tumors preserve IGM BCR expression.

**Figure 4.**
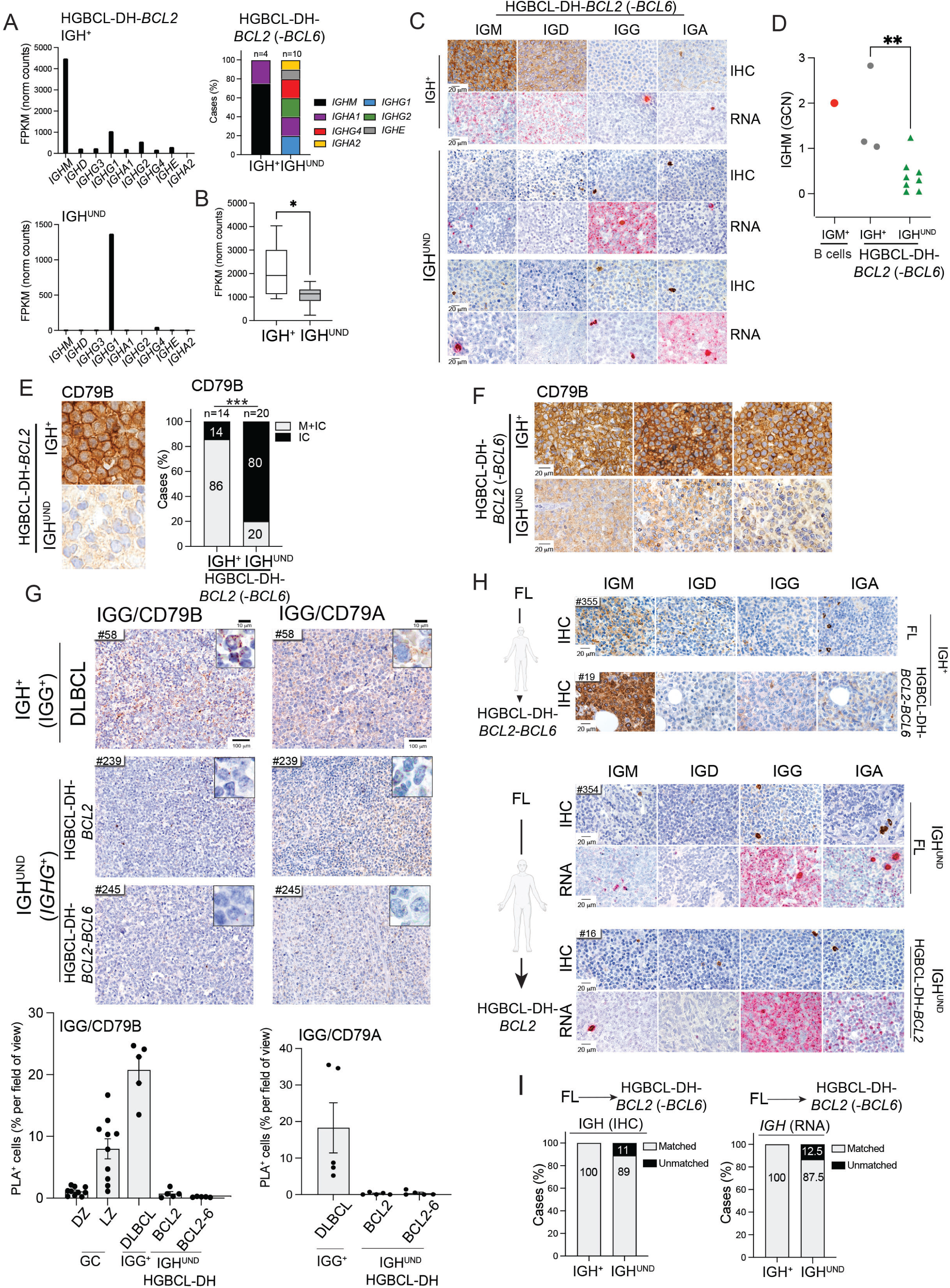
Class-associated IGH silencing in HGBCL-DH-*BCL2* is inherited from the FL precursor. (A) Class-specific *IGH* constant region transcripts in representative IGH^+^ (upper panel) and IGH^UND^ (lower panel) HGBCL-DH-BCL2 cases, quantified by RNA-seq. Histogram summarizes prevalent IGH transcripts in n=14 HGBCL-DH-*BCL2*. (B) Box plot representation of median and 5^th^-95^th^ percentile (whiskers) of most abundant *IGH* constant region transcript levels in IGH^+^ (white) and IGH^UND^ HGBCL-DH-*BCL2* (grey), measured by RNA-sequencing (*p≤ 0.05; unpaired t-test). (C) Class-specific IGH protein and transcripts in representative IGH^+^ (upper panel) and IGH^UND^ HGBCL-DH-*BCL2(-BCL6)* (mid and lower panels) cases detected by IHC and RNA scope, respectively. (D) Normalized *IGHM* gene copy number (GCN) in representative IGH^+^ (n=3; grey circles) and IGH^UND^ (n=8; green triangles) HGBCL-DH-*BCL2(-BCL6)*, measured by genomic qPCR. Purified IGM^+^ circulating B cells (n=1; red circle) controlled for two *IGHM* gene copies (**p≤ 0.01; unpaired t-test). (E) IHC localization of CD79B protein in representative IGH^+^ and IGH^UND^ HGBCL-DH-*BCL2*. Histograms summarize CD79B distribution scores in HGBCL-DH-*BCL2(-BCL6)* (n=34), discriminating intracellular (IC) from plasma membrane (M) immunoreactivity (***p≤ 0.001; Fisher’s exact test). (F) Reduced CD79B levels in representative IGH^UND^ HGBCL-DH-*BCL2(-BCL6)* cases (lower panels) as compared to IGH^+^ cases (upper panels), measured by IHC. (G) IGG/CD79B (left panels) or IGG/CD79A (right panels) protein complexes in an IGG- switched IGH^+^ DLBCL (upper panel) and in two representative IGH^UND^ HGBCL-DH-*BCL2(-BCL6)* (mid/lower panels), measured by PLA. Insets represent sections photographed at high magnification. Histograms depict mean frequency (+/- standard error of mean (SEM)) of PLA^+^ lymphoma cells in n=5 individual fields of view (black circles), quantified with HALO software. Quantification of CD79B/IGG complexes in GC DZ and LZ areas served as reference. (H) IGH IHC and in situ RNA analyses in representative IGH^+^ (top panel) and IGH^UND^ (mid/lower panels) FL-HGBCL-DH-*BCL2(-BCL6)* metachronous specimens. (I) IGH class correspondence between FL and metachronous/synchronous HGBCL-DH-*BCL2(-BCL6)* cases, assessed by either IHC (left) or RNA scope (right).

### IGH/CD79A/B complex shutdown and CD79B intracellular retention in IGH^UND^ HGBCL-DH-*BCL2*

In the endoplasmic reticulum of mature B cells, IGH and IG Light chains assemble with CD79A/B proteins to form the BCR complex, which gets transported to the cell surface after glycosylation of its subunits in the Golgi network. Given the marked IGH downregulation in IGH^UND^ HGBCL-DH-*BCL2*(*-BCL6*), we asked whether cellular distribution and expression of its interactor CD79B was affected in these tumors. In IGH^+^ HGBCL-DH-*BCL2*(*-BCL6*) (n=14) IGH expression was coupled to uniform plasma membrane associated CD79B immunoreactivity (Fig. 4E, Supplementary table S4D). In contrast, IGH^UND^ cases (n=20) exhibited preferential intracellular distribution (16/20) of CD79B, often associated with protein down-regulation (Fig. 4E-F, and Supplementary table S4D), resembling their GC DZ B cell precursors (Supplementary Fig. S4A-B). We performed in situ proximity ligation assays (PLA) to assess whether in IGH^UND^ HGBCL-DH-*BCL2*(*-BCL6*), CD79A and CD79B proteins could assemble into BCR complexes with, putative, residual IGH traces. IGH^UND^ HGBCL-DH-*BCL2*(*-BCL6*) expressing *IGHG* mRNAs, showed a significant reduction in IGG/CD79B or IGG/CD79A complexes (Fig. 4G and Supplementary table S4E). Conversely, IGG^+^ (or IGM^+^) DLBCL and GC LZ areas showed conspicuous IGG (or IGM)/CD79A/B PLA^+^ signals (Fig. 4G, S4C-D, and data not shown). IGH^UND^ tumors also showed marked contraction in the percentage of tumoral cells forming CD79A/CD79B heterodimers, as compared to IGH^+^ tumors (Supplementary Fig. S4E-F and Supplementary table S4E). Collectively, IGH silencing in HGBCL-DH-*BCL2*(*-BCL6*) withholds the assembly of IGH/CD79A/B BCR complexes, resulting in preferential intracellular distribution and downmodulation of CD79B protein.

### IGH silencing anticipates HGBCL-DH-*BCL2* onset

To assess timing of IGH silencing in HGBCL-DH-*BCL2(-BCL6)*, we analyzed three cases anticipated by a history of classical FL (17,18). Clonal relationship between metachronous tumors was established by sequencing *IGH::BCL2* and/or *BCL6::IGH* (in HGBCL-DH-*BCL2*-*BCL6*) translocation breakpoints, and *IGHV* rearrangements (Supplementary table S4F). In the IGH^+^ HGBCL-DH-*BCL2*-*BCL6*, IGM expression was shared with the preceding FL (Fig. 4H). Similarly, in two IGH^UND^ HGBCL-DH-*BCL2*, the IGH^UND^ phenotype and *IGHG* transcript pattern corresponded to those of the preceding FL (Fig. 4H). IGH matching was confirmed in10 additional FL-HGBCL-DH-*BCL2*(*-BCL6*) synchronous/metachronous pairs, except for a mixed IGM^+^/G^+^ FL, which evolved into an IGG-switched IGH^UND^ HGBCL-DH-*BCL2* (Fig. 4I, Supplementary table S4G). To assess whether metachronous HGBCL-DH-*BCL2*(-*BCL6*) cases arose through transformation of the preceding FL, or from a putative FL/ HGBCL-DH-*BCL2* common precursor cell (CPC) (58,59), we compared the mutational pattern of clonal *IGHV* rearrangements in the two consecutive tumors from three patients. In two patients diagnosed respectively with IGH^+^ HGBCL-DH-*BCL2-BCL6* (#19) and IGH^UND^ HGBCL-DH-*BCL2* (#297), the sharing of several *IGHV* mutations with the preceding FL was accompanied by acquisition of private nucleotide substitutions, suggestive of divergent evolution from a CPC (Supplementary Fig. S4G). The third patient suffered from an IGH^UND^ HGBCL-DH-*BCL2-BCL6* (#245), displaying a unique set of nucleotide substitutions which added up to those acquired by the preceding FL, compatible with linear progression of the disease. Together, these results postulate an early decision imposed on HGBCL-DH-*BCL2* precursor cells concerning IGH expression and class choice, implemented in the preceding FL or CPC.

### Bi-allelic *IGH* locus disruption in IGH^UND^ HGBCL-DH-*BCL2* infrequently intercepts *MYC*

Disruption of *IGH* alleles via interception in trans of the *BCL2* and *MYC* (and/or *BCL6*) loci through chromosomal translocations could interfere with normal *IGH* expression in HGBCL-DH-*BCL2*. To assess this, we conducted *IGH* break apart (BAP) DNA FISH analysis for 22 HGBCL-DH-*BCL2* (-*BCL6*) cases (Fig. 5A, S5A and Supplementary table S5). While most IGH^+^ HGBCL-DH-*BCL2*(-*BCL6*) (7/8; 87.5%) conserved one intact *IGH* locus, 43% (6/14) of IGH^UND^ tumors carried two disrupted *IGH* alleles (Fig. 5B and Supplementary table S5). *IGH::BCL2* dual-fusion FISH in IGH^UND^ HGBCL-DH-*BCL2*(-*BCL6*) with two disrupted *IGH* loci scored positive in every case (n=6; Fig. 5C and Supplementary table S5). Instead, *IGH::MYC* fusions scored positive by FISH only in 2/6 of these tumors (Fig. 5C and Supplementary table S5). All eight IGH^UND^ HGBCL-DH-*BCL2*(-*BCL6*) with single *IGH* locus breakage carried an *IGH::BCL2* fusion, including two cases also displaying an *IGH::MYC* rearrangement (25%; Supplementary table S5). IGH^+^ HGBCL-DH-*BCL2*(-*BCL6*) showed predominant monoallelic *IGH* locus disruption (7/8 cases; 87.5%), mostly (6/7 cases) consisting of *IGH::BCL2* fusions, whereas *IGH::MYC* rearrangements remained uncommon (2/8; 25%; Supplementary table S5). Collectively, in IGH^UND^ HGBCL-DH-*BCL2*(-*BCL6*), recurrent bi-allelic *IGH* chromosomal rearrangements infrequently intercept the *MYC* locus.

**Figure 5.**
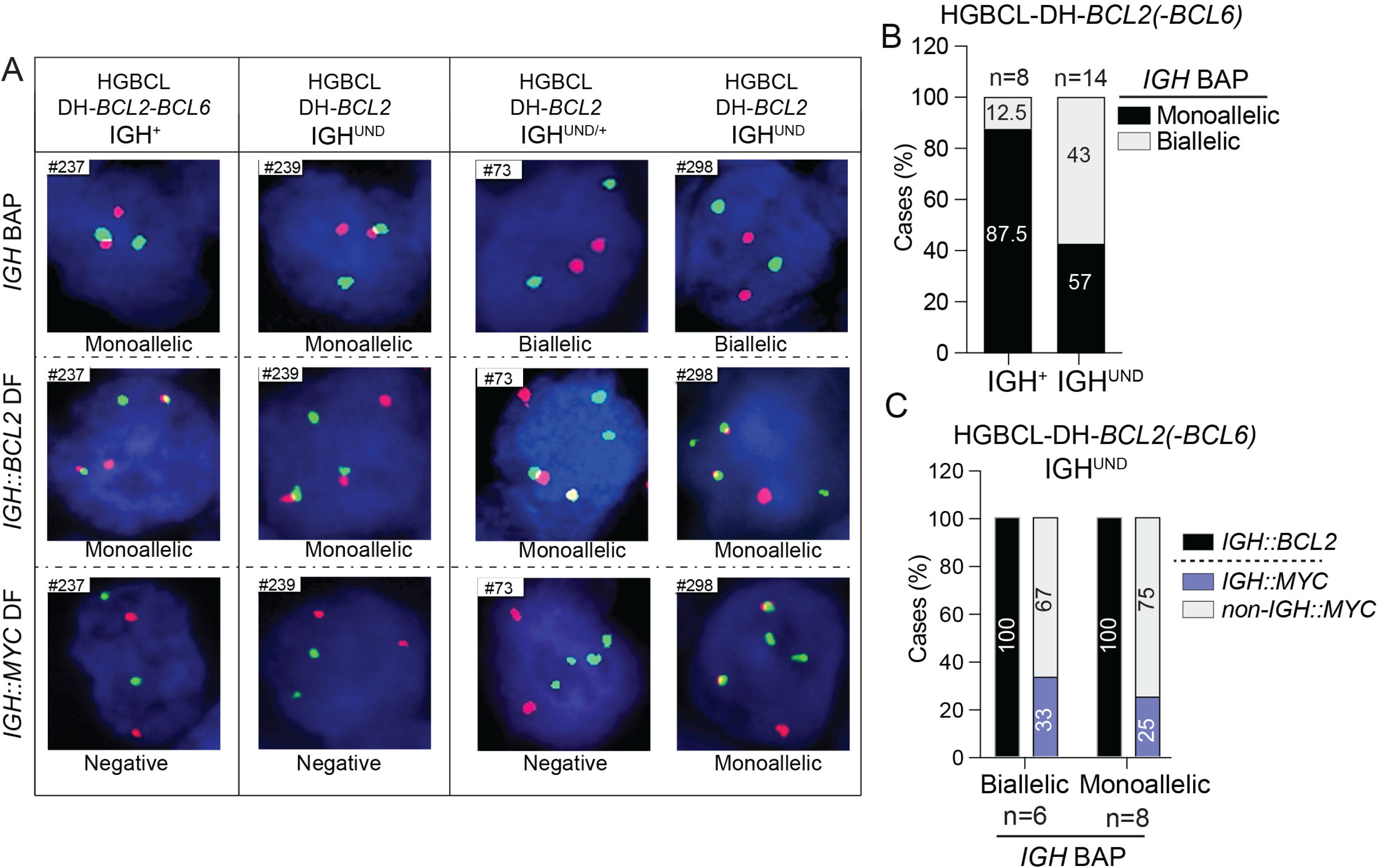
Bi-allelic *IGH* locus disruptions in IGH^UND^ HGBCL-DH-*BCL2* infrequently intercept the *MYC* gene. (A) Interphase DNA FISH analyses of representative IGH^+^ and IGH^UND^ HGBCL-DH-*BCL2(-BCL6)* to track *IGH* locus disruption (break apart FISH; BAP, top panel) and *IGH::BCL2* (mid panel) or *IGH::MYC* fusions (dual fusion or DF, middle/bottom panels). (B) Frequency of *IGH* mono/bi-allelic disruptions in IGH^+^ and IGH^UND^ HGBCL-DH-*BCL2* (*-BCL6*) cases (n=22). (C) Frequency of *IGH*::*BCL2* (black bar) and *IGH::MYC* (blue bar) fusions in representative IGH^UND^ HGBCL-DH-*BCL2(-BCL6)* cases with either one or two disrupted *IGH* loci. Cases with *non-IGH::MYC* rearrangements are represented as grey bars.

### RAG1/2 involvement in the progression from IGH^UND^ FL to HGBCL-DH-*BCL2*

The infrequent (∼ 25 %) occurrence of t(8;14) reported for HGBCL-DH-*BCL2* as compared to the high incidence detected in Burkitt lymphoma (> 70%) postulates a mechanism alternative to AID-induced DNA breaks at *IGH* switch regions, as causative of MYC rearrangements(60,61). To shed light on such process, we leveraged an IGH^UND^ FL/ HGBCL-DH-*BCL2*-*BCL6* metachronous pair (FL #353 and HGBCL-DH-*BCL2*-*BCL6* #245). Most FL cells presented an IGH^UND^ phenotype, with residual cells expressing IGG (Fig. 6A). RNA in situ analyses confirmed expression of *IGHG* transcripts replacing *IGHM*, consistent with bi-allelic IGG CSR (Fig. 6A). FISH analyses and whole genome sequencing (WGS) indicated bi-allelic disruption of *IGH* loci in FL cells due to *IGH::BCL2* and *IGH::BCL6* rearrangements occurring in trans (Fig. 6B and Supplementary Fig. S6A-B, Supplementary table S4F), predicting genetic extinction of IGH expression. Clonal relationship between the HGBCL-DH-*BCL2*-*BCL6* and preceding FL was supported by sharing the same *IGH/K V* gene rearrangements, *IGH::BCL2* and *IGH::BCL6* translocation breakpoints, IGH^UND^ phenotype and *IGHG* expression (Fig. 6A-B, Supplementary table S4F). WGS indicated acquisition by IGH^UND^ HGBCL-DH-*BCL2*-*BCL6* cells of a t(8;22)(q24;q11), confirmed by Sanger sequencing of the translocation breakpoint (Supplementary table S4F). The t(8;22) juxtaposed the *MYC* gene with the IG lambda (*IGL*) Joining-2 (*IGLJ2*) gene, at the recombination signal sequence (RSS). This result suggested the involvement of the RAG1/RAG2 recombinases in the translocation mechanism. In situ RNA scope and RNA-seq analyses revealed widespread *RAG1/2* expression in IGH^UND^ HGBCL-DH-*BCL2*-*BCL6* cells (Fig. 6C, Supplementary Fig. S6C, Supplementary table S6A). Screening of additional HGBCL-DH-*BCL2*(-*BCL6*) cases (n=34) identified *RAG1/2* expression in 26% (9/34), including both IGH^UND^ and IGH^+^ tumors (Fig. 6D, Supplementary table S6B). Similarly, *RAG1/2* transcripts were detected in GC-derived HGBCL-DH-*BCL2* (DOGUM, DoGKiT, WILL-2), DLBCL (SU-DHL-8, NU-DUL-1, TOLEDO) and BL (Namalwa) cell lines, including several ones positive for t(8;22)(q24;q11) (Supplementary Fig. S6D-E, Supplementary table S6C and (62)). Together, the results support a causal role for RAG1/2-mediated V-J recombination in promoting *IGL::MYC* rearrangements responsible for HGBCL-DH-*BCL2* transformation.

**Figure 6.**
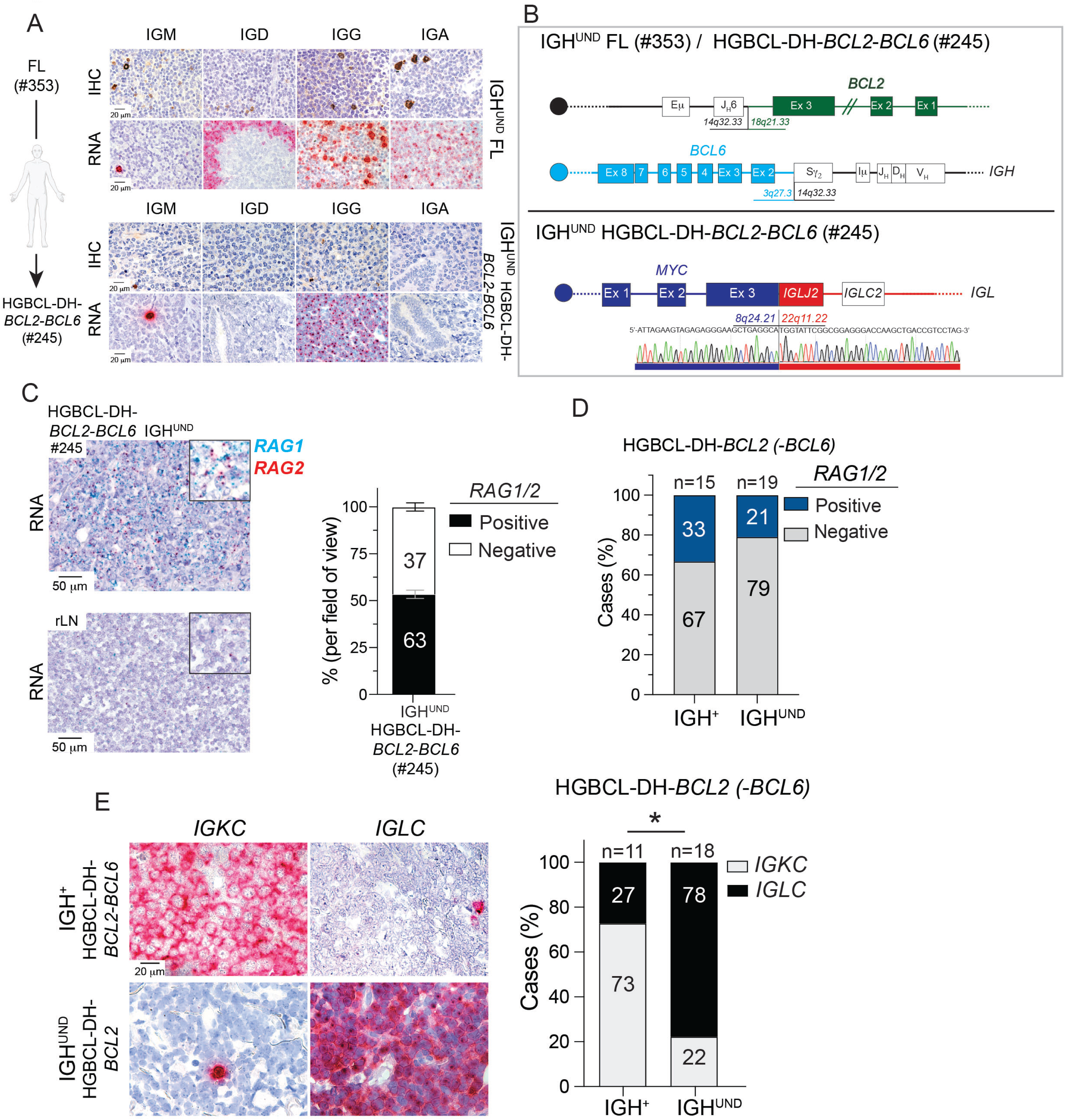
RAG1/2-driven progression of IGH^UND^ FL into HGBCL-DH-*BCL2*-*BCL6*. (A) IGH protein and RNA levels in metachronous IGH^UND^ IGG-switched, FL (#353) and HGBCL-DH-*BCL2*-*BCL6* (#245) specimens. In FL, *IGHD* transcripts lighten the mantle zone, while FL expression of *IGHG* and *IGHA* transcripts becomes restricted to *IGHG* upon progression to HGBCL-DH-*BCL2*-*BCL6*. (B) Structure of t(14;18) and t(3;14) chromosomal rearrangements in clonally-related FL (#353) and HGBCL-DH-*BCL2*-*BCL6* (#245) metachronous cases, as predicted by breakpoint sequencing. The t(8;22)(q24;q11) in HGBCL-DH-*BCL2*-*BCL6* juxtaposes *MYC* to *IGLJ2*. (C) In situ detection of *RAG1/2* transcripts in HGBCL-DH-*BCL2*-*BCL6* (#245) and in control rLN, measured by RNA scope. Histogram bars indicate mean frequency (+/- SEM) of *RAG1/2*^+^ cells measured in n=5 fields of view. (D) Summary of *RAG1/2* mRNA in situ detection in IGH^+^ and IGH^UND^ HGBCL-DH-*BCL2* cases (n=34), measured by RNA scope. Tumors with > 20% of RAG1/2-expressing cells were scored positive. (E) *IGKC* and *IGLC* transcripts in representative IGH^+^ and IGH^UND^ HGBCL-DH-*BCL2*, analyzed by RNA scope. Histograms summarize frequencies of *IGKC-* (grey) and *IGLC*- (black) expressing cases (n=29), among IGH^+^ and IGH^UND^ HGBCL-DH-*BCL2* cases.

### IG lambda light chain skewing in IGH^UND^ HGBCL-DH-*BCL2*

The juxtaposition of *MYC* to RSS sequences of *IGLJ2* segments detected in HGBCL-DH-*BCL2*-*BCL6* case #245 could reflect an attempt of the preceding FL to revise BCR specificity through IG light chain editing in response to BCR downregulation. Since BCR-edited healthy B cells predominantly express IG lambda light chains, we screened 36 HGBCL-DH-*BCL2*(-*BCL6*) for *IGKC/IGLC* transcripts in situ (Fig. 6E, Supplementary table S6D). Eight of 11 (73%) IGH^+^ HGBCL-DH-*BCL2*(-*BCL6*) were *IGKC^+^*, while remaining tumors transcribed *IGLC* mRNAs, in line with the ∼ 2:1 IGK/L ratio seen in healthy mature B cells (Fig. 6E and Supplementary table S6D). In sharp contrast, 78% of IGH^UND^ HGBCL-DH-*BCL2*(-*BCL6*) (14/18 cases) expressed *IGLC* transcripts (Fig. 6E and Supplementary table S6D). Seven additional tumors, including 5 IGH^UND^ HGBCL-DH-*BCL2*(-*BCL6*), expressed both *IGKC* and *IGLC* transcripts (Supplementary table S6D) and proteins (data not shown), indicating IG light chain isotype inclusion. Differential IG light chain isotype usage between IGH^+^ and IGH^UND^ HGBCL-DH-*BCL2*(-*BCL6*) was validated profiling representative cases for *IGK/LV* rearrangements using 5’RACE technology (Supplementary table S6E). In particular, in RAG1/2^+^ IGH^UND^ HGBCL-DH-*BCL2*-*BCL6* case #245, several productive and non-productive *IGKV* and *IGLV* rearrangements were captured in combination with a single, hypermutated, productive *IGHV* rearrangement, consistent with ongoing *IGV* light chain editing (Supplementary tables S6E and S3B) in the malignant clone. Similarly, several HGBCL-DH-*BCL2,* DLBCL and BL cell lines showed intraclonal IG light chain *V* gene diversification associated with *RAG1/2* expression (Supplementary table S6F). The latter tumor lines either displayed a sBCR/CD79B^UND^ phenotype, or preferentially expressed IGL^+^ BCRs (Supplementary table S6F, Fig. 7I, Supplementary Fig. S6E). In summary, GC-derived IGH^UND^ HGBCL-DH-*BCL2*(-*BCL6*) show signs of recurrent IG light chain V gene editing promoted by RAG1/2 expression.

**Figure 7.**
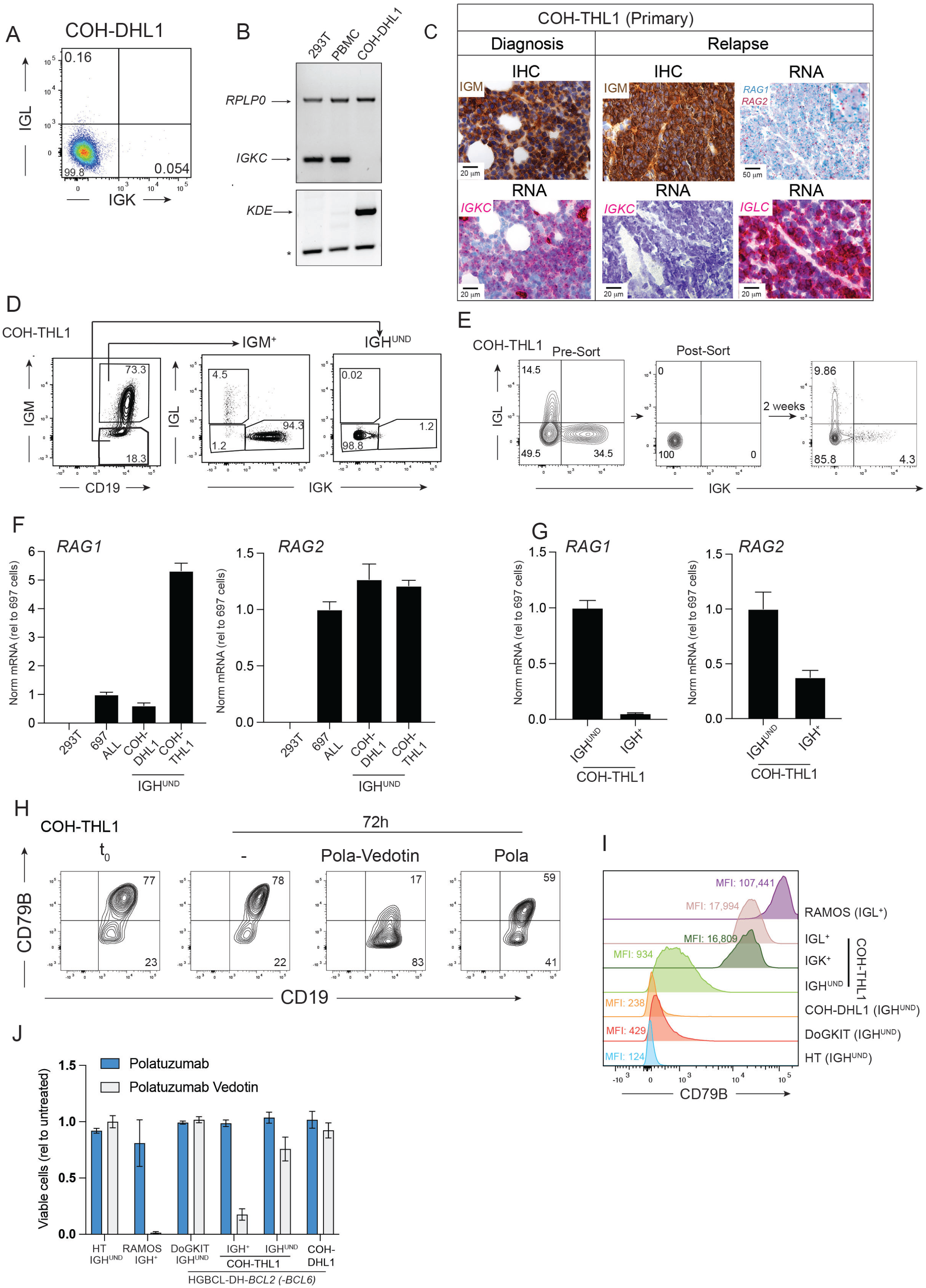
RAG-induced IG light chain editing in IGH^UND^ HGBCL-DH-*BCL2* cell line models influences response to Polatuzumab-Vedotin. (A) Representative (n=6) sIGK/IGL FACS analyses of COH-DHL1 cells. Cell frequencies within each quadrant are indicated. (B) PCR amplification of *IGKC* (upper panel) in COH-DHL1 cells, 293T cells and PBMC. Amplification of *RPLP0* controlled for DNA input. Genomic PCR for the *KDE* rearrangement was performed on the same sample set (lower panel). Data represent n=3 experiments. The asterisk indicates nonspecific bands. (C) IGM and IG light chain IHC and RNA analysis of the primary HGBCL-DH-*BCL2*-*BCL6* case and the COH-THL1 cell line derivative, analyzed at diagnosis (left panel) and after relapse (right panel). (D) Representative (n=6) FACS analysis of COH-THL1 cells before (left plot) and after (mid and right plots) gating on the indicated subsets. Numbers indicate cell frequencies (E). Representative (n=5) FACS analysis for sIGK/L of COH-THL1 bulk cultures (left panel), purified IGH^UND^ variants (mid panel) and derivatives analyzed 2 weeks later (right panel). Numbers indicate cell frequencies. (F) Normalized *RAG1/2* transcripts in IGH^UND^ COH-DHL1 and COH-THL1 bulk cultures relative to 697 leukemic cells, measured by qRT-PCR. 293T were included as negative controls. Data represent n=3 experiments. (G) Normalized *RAG1/2* transcripts in FACS-purified IGH^+^ and IGH^UND^ COH-THL1 cells, relative to 697 leukemic cells, quantified by RT-PCR. Data represent n=3 experiments. (H) Co-cultures of sCD79B^+^ (IGH^+^) and sCD79B^lo^ (IGH^UND^) COH-THL1 cells were treated for 72 h with Polatuzumab (Pola)-Vedotin, unconjugated Polatuzumab (Pola), or left untreated (-), and assessed by FACS for frequency of sCD79B^+^ cells. Data represent n=3 experiments. Numbers indicate cell frequencies in the corresponding quadrants. (I) Mean fluorescence intensity (MFI) values representing sCD79B levels quantified by flow cytometry for the indicated lymphoma cell lines. IGH^+^ COH-THL1 cells were separated into three subsets according to sIG light chain usage. (J) Cell viability measured 72 h post Polatuzumab-Vedotin/Polatuzumab treatment for the indicated DLBCL, BL and HGBCL-DH-*BCL2(-BCL6)* cell lines, measured by flow cytometry. Data are relative to untreated cells. Surface BCR status is indicated. Data represent n=3 experiments.

### IG light chain editing fuels HGBCL-DH-*BCL2* intratumor heterogeneity

To study the biological implications of RAG1/2 expression in HGBCL-DH-*BCL2*(-*BCL6*), we analyzed early-passage COH-DHL1 and COH-THL1 HGBCL-DH-*BCL2*(*-BCL6*) cell lines (19). COH-DHL1 cells carried the t(14;18)^+^ together with a non-*IGH::MYC* rearrangement (Supplementary Fig. S7A). Instead, COH-THL1 cells were positive for t(14;18)(q32.33;q21.3) and t(8;14)(q24.21;q32.33), with FISH analyses predictive of bi-allelic *IGH* locus disruption through rearrangements with *MYC* and *BCL2*. COH-THL1 cells carried an additional *BCL6* rearrangement intercepting the chr-8 derivative (der(8)t(8;14;3) (Supplementary Fig. S7A, B, (19)). COH-DHL1 cells showed undetectable surface IGH/K/L/CD79B, confirming and extending previous evidence (19) (Fig. 7A, 7I and Supplementary Fig. S7C). Similarly to IGH^UND^ HGBCL-DH-*BCL2*, COH-DHL1 cells expressed a single, in-frame, somatically mutated, isotype-switched (IGG1) *IGHV* rearrangement (Supplementary table S6G). COH-DHL1 cells inactivated both *IGKC* loci via RAG-dependent Kappa Deleting Element (*KDE*) rearrangements, followed by acquisition of an out-of-frame clonal *IGLV* rearrangement (Fig. 7B, Supplementary table S6G). The COH-THL1 cell line was established from the peripheral blood of a treatment-relapsed HGBCL-DH-*BCL2*-*BCL6* patient (19). The primary tumor evolved over time, shifting from IGK to IGL expression, which was linked to *RAG1/2* expression (Fig. 7C) co-occurring with IGM detection. COH-THL1 cells consisted of a mixture of CD79B^UND^/IGH^UND^ and CD79B^+^/IGM^+^ tumor B cells, with the latter grouped into IGK- or IGL-expressing cells (Fig. 7D, Fig. 7I). FACS-purified IGH^UND^ COH-THL1 cells reconstituted distinct pools of sIGK^+^ or sIGL^+^ tumor cells (Fig. 7E), unmasking the contribution of IGL chain editing to the IGH^UND^ to IGH^+^ phenotypic switch. BCR^+^ HGBCL-DH-*BCL2*-*BCL6* cells expressed several productive *IGKV* or *IGLV* rearrangements, while carrying a common somatically mutated in-frame *IGHV* rearrangement (Supplementary table S6G). Surface IGL^+^ HGBCL-DH-*BCL2*-*BCL6* cells mostly carried oligoclonal out-of-frame *IGKV* rearrangements, supporting IG light chain *V-J* recombination progressing from *IGK* to *IGL* loci (Supplementary table S6G). Conversely, IGH^UND^ HGBCL-DH-*BCL2*-*BCL6* cells transcribed a diverse set of out-of-frame *IGKV* and *IGLV* rearrangements, reconciling with the BCR-less phenotype. The same pool of cells also carried recently recombined in-frame *IGV* light chain rearrangements responsible for the IGH^UND^ to IGH^+^ phenotypic switch (Supplementary table S6G). IGH^UND^ COH-DHL1 and COH-THL1 cells expressed *RAG1/2* transcripts at levels comparable to, or higher than 697 pre-B leukemia cells serving as positive control (Fig. 7F). Importantly, *RAG1/2* expression was substantially downregulated in the IGH^UND^ to IGH^+^ switch, indicating a negative regulation imposed by sBCR on ongoing V-J recombination (Fig. 7G). Coherent with these results, inducible IGH extinction in primary λ-MYC; B1-8f murine B cell lymphomas significantly increased *Rag1/2* expression (Supplementary Fig. S7G), which, in turn, promoted BCR restoration in a fraction of the cells through secondary *IGHV* rearrangements in vivo (Supplementary Fig. S7E-J). Together, these results provide proof for ongoing IG light chain editing in human GC-derived HGBCL-DH-*BCL2*(-*BCL6*) and murine MYC-driven lymphoma models, contributing to bi-directional oscillation between sBCR^+^ and sBCR-less states and intra-tumor antigen receptor diversity.

### Reduced sensitivity of IGH^UND^ HGBCL-DH-*BCL2* cell line models to CD79B-targeted therapies

Preferential intracellular CD79B distribution associated to IGH silencing may hamper HGBCL-DH-*BCL2*(-*BCL6*) response to the anti-CD79B-drug conjugate (ADC) Polatuzumab-Vedotin (PV) recently approved for frontline treatment of DLBCL NOS and HGBCL(63). To test this, mixed cultures of IGH^+^ and IGH^UND^ COH-THL1 cells differing in sCD79B levels (Fig. 7H-I) were exposed for 72h to PV treatment. A single PV dose caused clearance of most sIGK/L/CD79B-expressing COH-THL1 cells, whereas IGH^UND^/CD79B^lo^ counterparts largely resisted the treatment, overtaking the culture (Fig. 7H). Reduced sensitivity reaching resistance to PV was confirmed in IGH^UND^ COH-THL1, COH-DHL1 and DoGKiT HGBCL-DH-*BCL2*-*BCL6* cell lines, behaving similarly to sCD79B negative HT DLBCL cells serving as positive control (Fig. 7I-J). In contrast, BL RAMOS cells, expressing the highest sCD79B levels among tested lymphoma lines, showed the highest sensitivity to the ADC. PV-induced lymphoma killing required BCR-dependent internalization of the payload, as DLBCL/BL/HGBCL-DH-*BCL2*(-*BCL6*) cell lines remained largely unaffected by treatment with unconjugated antibody, regardless of sCD79B status (Fig. 7J).

## Discussion

Comprehensive screening of primary DLBCL cases revealed approximately 35% exhibiting undetectable IGH immunoreactivity in most tumor cells. The failure to detect sIG light chains and IGH/CD79A/B complexes in representative cases indicated sBCR silencing in IGH^UND^ DLBCL. Preferential BCR shutdown in GCB-type DLBCL suggests a link between the tumor cell-of-origin and acquisition of the IGH^UND^ phenotype. IGH^UND^ lymphomas may inherit mechanisms of BCR downregulation from their GC B cell precursors (64–66), with transformation events trapping malignant B cells in a developmental state resilient to IGH silencing (16). Among GCB-type lymphomas with DLBCL morphology, HGBCL-DH-*BCL2*(*-BCL6*) exhibited IGH silencing in over 65% of the cases, consolidating and significantly extending previous scattered findings (61,67–70). Transcriptome and gene mutation profiling underscore the importance of BCR silencing in establishing a dark zone (DZ)-like molecular program characteristic of HGBCL-DH-*BCL2* (29,53,71). A subset of IGH^UND^ HGBCL-DH-*BCL2*(*-BCL6*) retained GC LZ-like transcriptional traits, possibly driven by BCR surrogate signals, or residual IGH/BCR, compatible with an origin from GC B cells arrested in transition between DZ and LZ state (72). In line with this, LZ-like IGH^UND^ HGBCL-DH-*BCL2(-BCL6)* showed an indeterminate DZ molecular signature (24) and closer transcriptional similarity to IGH^+^ cases. IGH^+^/BCR^+^ HGBCL-DH-*BCL2*(*-BCL6*) bypassed DZ-enforcing gene mutations, selecting others indicative of crosstalk with the TME, which featured richer T cell representation and immunosuppressive signals. Increased MYC expression and/or constitutive ERK/MAPK or PI3K signaling, predicted from mutational analyses, likely represent adaptive responses selected by IGH^UND^ HGBCL-DH-*BCL2*(*-BCL6*) cells to guarantee optimal growth under chronic BCR silencing(5). HGBCL-DH-*BCL2*(*-BCL6*) preserved potentially productive *IGHV* rearrangements, which remained refractory to structure-damaging mutations, while loosening signs of antigen-driven selection. These findings postulate the existence of a yet undefined pro-tumoral signal transmitted by the IGH chain to IGH^UND^ HGBCL-DH-*BCL2*(-*BCL6*) malignant B cells in a setting where IGH/CD79A/B and CD79A/B complexes remain below limit of detection. Predominant *IGHM* constant region gene expression in IGH^+^ tumors sharply contrasted with exclusive tracking of class-switched *IGH* constant region transcripts in the IGH^UND^ counterparts. IGH^+^ HGBCL-DH-*BCL2*(-*BCL6*) often displayed large deletions within *IGHM Sµ* switch regions (data not shown), suggestive of selection for enforced IGM expression (73) in a setting where persistent AID activity has favored CSR on the second, non-productive *IGH* locus carrying the *BCL2* rearrangement, in line with previous reports(57). In contrast, IGH^UND^ HGBCL-DH-*BCL2*(*-BCL6*) underwent bi-allelic IG CSR, indicating an origin from IGG^+^/A^+^ B cells. In healthy B cells, IGM BCRs strictly depend on the CD79 subunits for signaling, whereas IGG/A receptors bypass such requirement through own signaling-competent cytoplasmic domains (74–76). Surface BCR levels are differentially controlled between IGM^+^ and IGG^+^ GC B cells, with the IGG intracellular domain favoring stronger BCR internalization(77). These properties, together with lower transcript levels for BCR components in isotype-switched FL (7) and HGBCL-DH-*BCL2*(*-BCL6*) (our study), and weaker signals emanating from switched BCRs in lymphoma cells (78), could explain the peculiar propensity of IGG/A B cells to transform into IGH^UND^ DZ-like tumors. BCR shutdown in HGBCL-DH-*BCL2*(*-BCL6*) will also protect IGH class-switched malignant B cells from antigen-induced plasma cell differentiation (75,79,80). Consistent with this, we identified an uncommon isotype-switched IGA-expressing HGBCL-DH-*BCL2-BCL6* case, which selected a heterozygous *CD79A^Y188*^* truncating mutation expected to reduce BCR signal strength(4). Together, our data suggest that weak signals from IGH class-switched BCRs represent a permissive condition for t(14;18)^+^ GCB/(pre-)FL cell transformation into HGBCL-DH-*BCL2*, while, at the same time, hindering progression into DLBCL (78).

Our studies with metachronous pairs indicate that IGH levels and class choice are established prior to HGBCL-DH-*BCL2*(*-BCL6*) outgrowth, starting in the preceding FL, or in a CPC, including in situ follicular neoplasia, fueling both diseases (58,59,81). IGM^+^ FL cells (or CPC) recurrently select de novo acquired N-linked glycosylation sites within IGHV domains, introduced by *IGV* somatic hypermutation (SHM) (82–84). Oligomannose-rich FL BCRs engage with extracellular host or bacterial mannose-binding lectins to deliver weak antigen-independent signals sustaining the growth of malignant B cells (85–88). This setting may select for persistent sBCR expression in IGM^+^ FL/CPC (84,86), transmitting such dependence to the HGBCL-DH-*BCL2*(*-BCL6*) derivative. Instead, IGH-switched FL cases express autoreactive BCRs more often than their IGM^+^ counterparts, possibly promoting chronic BCR downregulation/desensitization as adaptive response to sustained signaling (86,89,90). Escape from self-reactivity has been also proposed for IGG DLBCL(91). Concomitant selection in IGH-switched FL of de novo N-glycosylation sites in IGHV CDR/FWR could favor engagement of residual IGH molecules with intracellular lectins triggering a low-strength survival signal, reconciling BCR silencing with *IGHV* FWR negative selection for R mutations observed in HGBCL-DH-BCL2 derivatives (89,92). Such dynamics may impose the progressive selection in FL for gene mutations enforcing an IGH^low/UND^ GC DZ-like state, ultimately driving transformation into IGH^UND^ HGBCL-DH-*BCL2*(*-BCL6*). Similarly to BCR- edited autoreactive B cells (93), IG-switched IGH^UND^ HGBCL-DH-*BCL2*(*-BCL6*) showed strong IGL light chain skewing, while IGM^+^ cases preferentially expressed IGK chains. We postulated that chronic BCR downregulation in CPC/FL/HGBCL-DH-*BCL2*(*-BCL6*) triggered RAG-induced IG light chain revision (89,94–96). We confirmed *RAG1/2* expression in 26% of HGBCL-DH-*BCL2*(*-BCL6*) cases. Also, performing longitudinal *IGK/LV* gene tracking, we provide experimental proof for ongoing RAG-driven IG light chain editing in a primary HGBCL-DH-*BCL2*-*BCL6* case extending analyses to the COH-THL1 cell line derivative(19), reconciling previous observations (97–101). COH-THL1 cultures spontaneously generated marked intra-tumor BCR diversity consisting of sIGK^+^, sIGL^+^, sIGK^+^/L^+^ double-producer tumoral B cells co-existing with a population of IGH^UND^ cells. HGBCL-DH-*BCL2*-*BCL6* intra-tumor diversification through *IGV* light chain editing was confirmed in an IGH^UND^ case (#245) evolving from FL, and in several HGBCL-DH-*BCL2* and DLBCL cell lines, which either lost, or primarily expressed IGL-restricted BCRs. Whereas COH-THL1 cells allowed live monitoring of an IGH^+^-to-IGH^UND^ phenotypic switch in HGBCL-DH-*BCL2*-*BCL6*, the COH-DHL1 tumor line exemplifies the dead-end of an evolutionary trajectory which led a GC-derived IGG1/K^+^ HGBCL-DH-*BCL2* to irreversibly lose BCR expression through consecutive rounds of unproductive RAG-dependent *IGK/L*V rearrangements, while preserving *IGHV* productivity. Notably, IGH^UND^ COH-THL1 cells occasionally restored sBCR expression through secondary productive *IGV* light chain rearrangements. These observations indicate that during HGBCL-DH-*BCL2* evolution, malignant cells may bi-directionally oscillate between IGH^+^ and IGH^UND^ states, fueled by repeated rounds of *IGV* light chain revision. IGH^UND/+^ mixed cases identified in our initial IHC screen may capture such dynamics. Studies with the λ-MYC murine lymphoma model, and human IGH^UND^ HGBCL-DH-*BCL2* cell lines, supported by previous work on immature and GC B cells (102–104), and DLBCL (94) cells, anticipate that silencing/weakening of BCR signaling may favor *RAG1/2* re-expression in GC-derived lymphomas. This scenario is more likely to happen in IGH^UND^ tumors but could also occur in IGH^+^ tumors especially those experiencing weak BCR signals, as indicated by concomitant IGM and *RAG1/2* expression in a subset of HGBCL-DH-*BCL2* primary cases and cell lines. Tumor cell-intrinsic genetic mutations and/or microenvironmental cues (i.e. cytokines) may also contribute to *RAG1/2* deregulation in HGBCL-DH-*BCL2*(*-BCL6*), with the BCR opposing such action proportionally to its signal strength, as witnessed in COH-THL1 cells.

The low frequency of t(8;14) translocations reported for HGBCL-DH-*BCL2*(*-BCL6*)(60,61), and confirmed by our study, could be interpreted with the need of malignant B cells to preserve the functionality of the productive *IGH* locus, once the second allele is functionally disrupted by the t(14;18). However, t(8;14) remained infrequent even in IGH^UND^ HGBCL-DH-*BCL2*(*-BCL6*) cases with two disrupted *IGH* loci, calling for mechanisms other than aberrant IG CSR/SHM drifting *MYC* translocations away from *IGH*. A case study based on IGH^UND^ FL-HGBCL-DH-*BCL2*-*BCL6* metachronous specimens (#353/#245) allowed us to link acquisition of t(8;22)(q24;q11) with *RAG1/2* recombinase activity. Sequencing the translocation breakpoint revealed the juxtaposition of *MYC* to *IGJL2*, surgically replacing the 5’ RSS sequence, consistent with illegitimate V-J recombination. The capture of multiple *IGV* light chain rearrangements confirmed ongoing RAG1/2-mediated recombination in this tumor. The causal relation between RAG1/2 expression and *IGL::MYC* translocations in isotype-switched GC-experienced IGH^UND^ B cell lymphomas has been reported in conditional knock-out mice for the DNA double-strand (ds) break repair protein XRCC4 in *Tp53*-deficient mature B cells (105,106). Our data, which are fully consistent with the latter observations, support a model whereby IGH^UND^ HGBCL-DH-*BCL2*(*-BCL6*), originating from IGH-switched (pre-)FL B cells, activate *IGLV* gene editing in response to BCR downregulation. Under such circumstances, RAG1/2 (re)expression will expose *IGLV* loci to DNA ds breaks at RSS sites, which may intercept broken *MYC* loci, favoring t(8;22). In line with this model, recurrent acquisition of t(8;22) has been repeatedly reported in aggressive B cell lymphomas with *MYC* and *BCL2* rearrangements, including IGH^UND^ HGBCL-DH-*BCL2* evolving from FL (61,70,107–114). RAG-induced DNA breaks at cryptic RSS may also contribute to *non-IG*::*MYC* rearrangements in HGBCL-DH-*BCL2*, especially in a setting of impaired DNA repair (115). The expression of *RAG1/2* in cell lines representative of BL and GCB-DLBCL predicts a broader contribution of the recombinases to the acquisition of structural variants and, more broadly to the fueling of genome instability in MYC-driven B cell lymphomas, supported by previous observations (116,117).

Recent meta-analyses of B-NHL clinical studies have highlighted lower efficacy of therapies based on Polatuzumab-Vedotin in GCB-type DLBCL and HGBCL-DH-*BCL2* (116). Our data showing recurrent IGH silencing in GCB-DLBCL and HGBCL-DH-*BCL2* cases, leading to preferential intracellular CD79B retention, supported by limited sensitivity of IGH^UND^ GCB DLBCL and HGBCL-DH-*BCL2* cell line models to PV, offers an explanation for the clinical results.

A solid body of evidence has consolidated the notion that BCR signaling contributes to onset and persistence of several mature B cell neoplasms, rendering it an effective target of therapy (3). Here, we present first-time evidence that chronic BCR silencing shapes the evolution of HGBCL-DH-*BCL2* from (pre-)FL and contributes to its onset. Our findings urge for careful consideration of sCD79B-targeted regimens for the cure of HGBCL-DH-*BCL2*, and more broadly GCB-type DLBCL NOS, emphasizing routine monitoring of surface IGH expression, or biomarkers thereof, for treatment indication.

### Limitations of the study

HGBCL-DH-*BCL2*(*-BCL6*) represent infrequent B cell malignancies rendering biological studies challenging and therefore identification of improved therapies an unmet medical need. We tackled this, collecting through a multi-center study nearly 100 HGBCL-DH-*BCL2*(*-BCL6*) specimens, ensuring the largest study on BCR biology on this lymphoma subtype to date. Despite the limited number of HGBCL-DH-*BCL2*(*-BCL6*) cases clustered for IGH expression and profiled for bulk and spatial transcriptomics, we extrapolated information on tumor-infiltrating T cells, which we successfully validated by IHC analyses in a largely independent cohort. Lack of reference germline DNA limited the power of whole exome analyses yet allowed individual HGBCL-DH-*BCL2*(*-BCL6*) cases to be ontogenetically related to putative DZ/LZ B cell precursors, which was independently validated by spatial and bulk transcriptomics. The initial analysis of a FL/ HGBCL-DH-*BCL2*-*BCL6* metachronous pair revealing a link between BCR silencing and *RAG1/2* expression, found confirmation in additional HGBCL-DH-*BCL2*(*-BCL6*) cases, seeking mechanistic insights in multiple patient-derived and mouse lymphoma models. The clinical impact of the IGH^UND^ phenotype on HGBCL-DH-*BCL2* and DLBCL patient response to CD79B-targeted therapies and to more widely used glucocorticoid-based chemo-immuno regimens (i.e. R-CHOP) impacting on BCR signaling (117), awaits controlled trials.

## Authors’ contributions

**P. Sindaco**: Data curation, formal analysis, validation, investigation, visualization, methodology, writing-original draft. **S. Lonardi**: Data curation, formal analysis, validation, investigation, visualization, methodology, writing-original draft. **G. Varano**: Data curation, formal analysis, validation, investigation, visualization, methodology, writing-original draft. **I. Pietrini**: formal analysis, validation, investigation, visualization, methodology. **G. Morello**: Formal analysis, validation, investigation, visualization, methodology. **P. Balzarini**: Formal analysis, validation, investigation, visualization, methodology. **H. Neuman**: Software, formal analysis, investigation, visualization data curation. **F. Vit:** Software, formal analysis, investigation data curation. **G. Bertolazzi**: Software, formal analysis, data curation, **H. Arima**: Investigation. **M. Chiarini**: formal analysis, investigation, **L. Lorenzi**: Formal analysis, investigation. **V. Pellegrini**: Investigation, visualization. **S. Giampaolo**: Investigation, formal analysis. **D. Garzon**: Validation. **C. Ranise**: Investigation, formal analysis. **M. Bugatti**: Investigation, methodology. **C. Pagani**: Investigation. **R. Daffini**: Investigation. **F. Mainoldi**: Investigation. **V. Selvarasa**: Investigation. **A. Sivacegaram**: Investigation. **H. Yang**: Supervision. **L. Ying**: Formal analysis, validation, investigation, data curation. **V. Cancila**: Investigation. **R. Bonnal**: Formal analysis. **E. Visco**: Investigation. **C. Lopez Gonzalez**: Visualization, **P. Capaccio**: Resources, **A.J.M. Ferreri**: Resources **A. Tucci**: Resources, **A.D. Cabras**: Resources, **G. Pruneri**: Resources, **A. Di Napoli**: Resources, **R. Siebert**: Supervision, **R. Bomben**: Supervision, formal analysis, investigation. **B. Falini**: Resources, **M. Pizzi**: Resources, formal analysis, investigation. **J.Y. Song**: Resources. **W.C. Chan**: Resources. **M. Ponzoni**: Supervision, formal analysis, investigation. **R. Mehr**: Supervision, formal analysis, investigation, writing original-draft, **C. Tripodo**: Supervision, formal analysis, investigation, methodology. F. Facchetti: Supervision, resources, formal analysis, visualization, methodology, project administration, writing-review and editing. **S. Casola**: Conceptualization, supervision, resources, formal analysis, visualization, methodology, project administration, writing-review and editing.

## Acknowledgments

This work was supported by grants from the Italian Association for Cancer Research (AIRC; IG #23747 to S.C. and #22145 to C.T.), the Italian Ministry of Education, University and Research (PRIN #20175L9H7H to F.F., M.Po., M.Pi., and PRIN #2017K7FSYB to C.T.) and the US-Israel Binational Science Foundation grant 2013432 (to RM). P.S. was supported by post-doctoral fellowships sponsored by AIRC AND PRIN #20175L9H7H. S.L and L.L. were supported by Fondazione Beretta per la Ricerca sul Cancro. G.V received support from an American Society of Hematology Global Research Award (AGRA2023-4) and the Marie Skłodowska-Curie post-doctoral training program (H2020-MSCA-IF-2019 #895887). S. G. is supported by a Marie Skłodowska-Curie postdoctoral fellowship (H2022-MSCA-IF-2022-PF-01 # 101111183) and the CARIPLO Foundation (Progetto Giovani Ricercatori). D.G. received support from the Marie Skłodowska-Curie Innovative Training Network (H2020-MSCA-ITN-765158-COSMIC). H.N. was supported by a Bar-Ilan University President’s scholarship. We thank the IFOM Imaging (M.G. Totaro and D. Parazzoli) and Cellular & Preclinical Models (I. Rancati, S. Lavore, G. Ossolengo) technological development units, and the COGENTECH Genomics (S. Minardi and M.Riboni) and Histopathology (F. Pisati) units, for support in flow cytometry, cell sorting, cell line biobanking and molecular profiling, NGS and histopathology. Thanks to Dr. A. Tironi and Dr. M. Ungari for contributing biopsy specimens, and S. Zini, A. Valzelli, M. Tomaselli, P.Bossini, A. de Zorzi, L. Fappani, L. Fontana, T. Gulotta, S. Rosola, E. Albertini, M. Benedetti, F. Filippini and L. Zavaglio for technical and administrative support.

## Materials and Methods

### Human participants

All samples in this study were collected with informed consent for research use from patients enrolled on clinical protocols approved by Institutional ethical committee boards (ASST Spedali Civili Brescia approval n. NP3719-FF-BCR2, Fondazione IRCCS Istituto Nazionale Tumori n.approval INT 218/21, Fondazione IRCCS Ca’ Granda Ospedale Maggiore Policlinico, Milano approval n. 707_2020), complying with ethical regulations and in accordance with the Declaration of Helsinki, at the respective collection centers: Azienda Socio-Sanitaria Territoriale degli Spedali Civili di Brescia (Italy), San Raffaele Hospital (Milan, Italy), Istituto Nazionale Tumori Milano (Italy), Azienda Ospedaliero Universitaria Padova (Italy). A total of 352 patients with a diagnosis of Diffuse Large B cell Lymphoma (DLBCL, including Germinal-Center [GCB] or non-Germinal-Center B cell [non-GCB] subtypes, or High-Grade B cell Lymphomas (HGBCL) with *MYC* and *BCL2* rearrangements (with and without BCL6 rearrangements) (“double-hit”, HGBCL-DH-*BCL2*)(17,18) were included in the study. Two main cohorts of patients were enrolled. The first consisted of a total of n= 253 consecutive DLBCL and HGBCL-DH-*BCL2*(-*BCL6*) cases (Table S1A), including: 1) n= 156 formalin-fixed paraffin-embedded (FFPE) tumor biopsies (# 1-156, Supplementary table S1A) selected from n=563 consecutive DLBCL and HGBCL patients diagnosed between January 2016 and December 2020, at the Pathology unit of Spedali Civili, Brescia (Italy); 2) n= 75 tissue microarray (TMA) biopsies (# 157-231) from 372 consecutive DLBCL and HGBCL cases diagnosed between January 2011 and December 2016 at Azienda Ospedaliero-Universitaria di Padova (Italy); 3) n= 22 consecutive DLBCL and HGBCL (# 331-352) specimens prospectively collected between October 2020 and December 2023 at Spedali Civili, Brescia (Italy). Low-quality samples with extensive fibrosis and/or necrosis, and/or with insufficient material (i.e., biopsies smaller than 15 mm) were excluded from analyses. Cases with both nodal (n=172) or extranodal (n=81) biopsies were included in the study. The second cohort was restricted to GCB MYC and BCL2 dual expressor lymphomas (MB2 DE) only, comprising n= 56 MB2 DE non-HGBCL-DH-*BCL2(-BCL6)* and HGBCL-DH-*BCL2(-BCL6)* from the consecutive cohort, and n= 97 additional cases (# 232-328, Supplementary table S1B), the latter obtained through a multicenter collection from Spedali Civili, Brescia (Italy), San Raffaele Hospital, Milan (Italy) and Istituto Nazionale Tumori, Milan (Italy). Two additional HGBCL-DH-*BCL2(-BCL6)* cases were kindly provided by dr. Chan (City of Hope National Medical Center, United States). A total of n= 93 HGBCL-DH-*BCL2(-BCL6)* cases were included in this cohort. Reactive tonsil and lymph node samples from healthy donors were used as controls.

### Pathology review

The definition of “cell of origin” (COO) was established according to Hans’ algorithm (CD10, BCL6, MUM-1 expression), together with MYC and BCL2 protein expression (cut off for positivity, respectively ≥40% and ≥50%). IGH immunostainings were centralized and systematically evaluated by two pathologists (FF, SL), selecting tissue areas devoid of diffuse fibrous tissue, numerous apoptotic cells, diffuse necrosis, or with diffuse intercellular tissue stain. As further confirmation, immunostained digitalized images (Aperio Scanscope CS from Leica Microsystems), were evaluated by three additional pathologists (San Raffaele Hospital, Milan, Spedali Civili, Brescia, University of Padova, Padova), reaching a concordance of 0.99 (Cohen’s Kappa). The IGM stain was performed on TMA samples in two different laboratories (Spedali Civili, Brescia, and University of Padova), with a concordance for IGM detection of 0.97. Any non-concordant diagnoses among pathologists were re-reviewed, and resolution was achieved by performing independent immunostainings in a second center.

### Immunohistochemical analysis

FFPE tissue sections were immunostained for IGH chains IGA, IGD, IGG, and IGM, CD79B, CD20, and CD3 using the Bond Max/Bond-III (Leica Microsystems) or the BenchMark Ultra (Ventana Medical Systems) autostainers (antibodies listed in the Table 7). AICDA and CD79B (Abcam) stainings were performed manually. Deparaffinized slides were subjected to antigen unmasking procedure using a water bath at 98°C for 40’ in buffer Tris-EDTA pH 9.0 or Citrate pH 6.0 and then incubated for 1h at room temperature with the antibody diluted 1:1000 and 1:250, respectively. Novolink Polymer was used as a detection system, and chromogen reaction was developed using 3,3’Diaminobenzidine (DAB). Immunostained sections were photographed with an Olympus DP-73 WDR digital camera mounted on Olympus BX60 microscope. CellSens standard 1.17 was used as acquisition software. Digitalization through Aperio ScanScope CS (Leica Microsystems) was also used to take images at low magnification. Positivity was determined by the presence of complete membranous and/or cytoplasmic reactivity in tumor cells. Cases were classified according to the percentage of IGH-expressing cells. Specifically: cases with over 90% of tumor cells exhibiting IGH immunoreactivity were categorized as IGH-positive (IGH^+^). Conversely, when IGH reactivity was undetectable in more than 90% of cells, the tumor was classified as IGH-undetectable (IGH^UND^). Cases showing IGH expression in a range between 10% and 90% of atypical cells were classified as IgH^UND/+^. Plasma cells acted as internal IGH staining control. AID stain was scored in HGBCL-DH-*BCL2*(-*BCL6*) cases (n=26) and defined as positive when reactivity was detected in over 25% of tumor cells. CD79B expression in HGBCL-DH-*BCL2*(-*BCL6*) (n=34) was scored as cytoplasmic vs membrane-bound, regardless of intensity. Quantification (number of cells/mm^2^) of CD3^+^ infiltrating lymphocytes was performed on digitalized slides from IGH^+^ and IGH^UND^ DHL/THL cases (n=55), using the Aperio IHC Nuclear Image analysis algorithm in ImageScope v.12.3.2.8013 (Leica Microsystems). Cell counting was performed in 1 mm^2^ tumor-rich areas.

### B cell lymphoma and leukemia cell lines

Human B cell lymphoma cell lines SU-DHL-4 (M, 38 Years (Y)), SU-DHL-6 (M, 43 Y), SU-DHL-8 (M, 59 Y), DOGKIT (M, 56 Y), WILL-2 (F, 63 Y), NU-DUL-1 (M, 43 Y), NU-DHL-1 (M, 73 Y), NAMALWA (M, 3 Y), 697 (M, 12 Y), DOGUM (F, 49 Y), HT (M, 70 Y), HeLa (F, 31 Y), and Ramos (M, 3 Y) were purchased from DMSZ and cultured according to biorepository instructions in RPMI-1640 supplemented with 10% heat-inactivated fetal bovine serum (FBS) (Sigma-Aldrich) and 2 mM L-glutamine (Euroclone). COH-DHL1 (M, 58 Y) and COH-THL1 (M, 78 Y) cell lines(19) were kindly provided by Dr. Chan and cultured in RPMI-1640 medium (Euroclone) supplemented with 10% heat-inactivated FBS (Sigma-Aldrich), 2 mM L-glutamine and 10mM HEPES (Euroclone), at a density of 0.5 - 1.5 x 10^6^ cells/ml. λ-MYC;B1-8f mouse lymphoma lines(5) were cultured at 2× 10^5^ to 2.5 × 10^5^ cells/mL in high-glucose DMEM supplemented with 10% heat-inactivated FBS, 0.1mM nonessential amino acids, 1mM sodium pyruvate, 50μM β-mercaptoethanol (Thermo Fisher Scientific) and 2mM L-glutamine (Euroclone). Cell lines were periodically tested for Mycoplasma (MycoAlert mycoplasma detection kit, Lonza) and grown at 37°C in a humidified atmosphere with 5% CO2. Cell line authentication was performed using the GenePrint 10 System (10-Locus STR System for Cell Line Authentication, Promega).

### Cytoblock preparation

SU-DHL-4 and SU-DHL-6 lymphoma cell lines acted as IGH class-specificity controls for IHC stainings (Figure S1C). Pre-warmed (40°C) low melting 1% agarose (Euroclone GellyPhor LM) solution (in PBS) was mixed with 1 x 10^7^ lymphoma cell suspensions and solidified in a conical stamp for at least 30 minutes. Following agarose embedding, blocks were removed from the stamp, fixed with 4% paraformaldehyde for 4 hours, washed with 70% ethanol, and embedded as paraffin blocks. The same procedure was used to establish IGH/L status in COH-DHL1 and COH-THL1 HGBCL-DH-*BCL2*(-*BCL6*) cell line models.

### Generation of anti-human CD79B monoclonal antibody

Anti-human CD79B mouse monoclonal antibody (clone # 128-4C5) was established by B.F. at the OncoHematological Research Center (CREO), Perugia University. The mAb was raised against a recombinant peptide corresponding to the CD79B protein’s amino terminal extracellular portion. Specificity was confirmed by immunoblotting and flow cytometry using CD79B-positive (normal B cells and transformed B cell lines SU-DHL16 and RIVA) and negative (T lymphocytes and the acute myeloid leukemia cell line KG-1) cells, matching results with those obtained with a commercial anti-CD79B antibody. Immunoreactivity against 293T cells complemented with CD79B cDNA or flow cytometric and immunoblotting analyses on *CD79B* gene-edited/knock-out SU-DHL-16 cells were included as additional 128 4C5 specificity tests. Anti-CD79B staining on FFPE tissue sections was performed on BOND-MAX autostainer (Leica Biosystems) using the EDTA 15 antigen retrieval protocol and incubating the antibody (dilution 1:4) for 30 minutes.

### Human primary B-lymphoma cell suspensions

Primary lymphoma cell suspensions were prepared from freshly collected nodal biopsies. After surgical resection, tumor specimens were placed in isotonic PBS buffer. A 5-10 mm^3^ tissue fragment was transferred in dissociation medium (RPMI-1640 supplemented with L-glutamine and penicillin/streptomycin), finely minced and subjected to tissue dissociation into single-cell suspension using Gentle MACS™ dissociator (Miltenyi Biotec). Cell suspensions were filtered through a 70-µm cell strainer to remove aggregates and centrifuged at 1200 RPM for 5 minutes at 4°C. Residual red blood cells (RBC) were lysed incubating cell suspension for 3 minutes at 4°C with 1X RBC Lysis Buffer (BioLegend). Cell suspensions were washed with PBS, counted, and used for downstream analysis or cryopreserved in freezing solution (90% FBS, 10% dimethylsulfoxide) in liquid nitrogen.

### Flow cytometry

Between 0.5 and 1×10^6^ primary B lymphoma cells or B lymphoma lines were washed in PBS and stained in FACS Buffer (1% BSA, 2mM EDTA buffer, 0.01% sodium azide, in PBS) with combinations of fluorescently labeled antibodies (Listed in the Supplementary table S7) for 20 minutes in the dark at 4°C. Stained cells were washed twice with FACS buffer and analyzed on BD FACSCanto II cell analyzer (BD Biosciences) or on a Cytek Aurora spectral analyzer (Cytek Biosciences). Alternatively, cells were purified in sorting medium (DMEM supplemented with 30% FBS, 2 mM L-glutamine, 1X Penicillin/Streptomycin solution) using a FACS Aria Cell sorter (BD Biosciences). Data analysis was performed using FACSDiva (BD Biosciences), SpectroFlo (Cytek Biosciences), and FlowJo v. 10.9.0 (BD Biosciences) softwares.

### Fluorescence in situ hybridization

Fluorescence in situ hybridization analysis (FISH) was performed on FFPE tissue sections using the IQFISH break apart probes for *MYC* (8q24), *BCL2* (18q21), *BCL6* (3q27)(Agilent Technologies) and *IGH* (14q32.33) loci (ZytoLight SPEC IGH Dual Color Break Apart Probe from ZytoVision); *IGH-MYC*/CPP8 Tri-color Fusion/Translocation FISH Probe Kit (CytoTest) and ZytoLight SPEC *BCL2/IGH* Dual Color Dual Fusion Probe (ZytoVision) were used to identify the translocation partner of *IGH*. FlexISH *BCL2/BCL6* DistinguISH Probe (ZytoVision) was employed to check if *BCL2* and *BCL6* rearrangements involved the same chromosome. FISH was carried out according to manufacturer’s guidelines. At least n= 100 cells were evaluated, and FISH images were captured at x 100 magnification using the Nikon Eclipse 90i microscope and elaborated using the Genikon software 3.7.17 (Nikon), or at x 63 magnification using Leica DM6000B Automated Microscope and elaborated using CytoVision MB8 software (Leica Microsystems).

### RNA in situ hybridization

RNA in situ hybridization was performed on FFPE sections, using Advanced Cell Diagnostics probes complementary to *IGHA1*, *IGHD*, *IGHM*, and the pools of *IGHG*, as well as *IGKC* and *IGLC*, following manufacturer’s instructions. Probes complementary to *UBC* and *DAPB* were used to control for RNA quality before staining with target-specific probes. Briefly, freshly cut 4-μm FFPE sections were deparaffinized in xylene, treated with RNAscope hydrogen peroxide solution (Advanced Cell Diagnostics) for 10 minutes at room temperature, subjected to retrieval for 15 min at 98°C in 1x Target Retrieval reagent (Advanced Cell Diagnostics) and protease treatment with RNAscope Protease Plus (Advanced Cell Diagnostics) at 40 °C for 30 minutes. Hybridization was performed for 2 hours at 40°C, and signal was revealed using RNAscope 2.5 HD Detection Reagent FAST RED (Advanced Cell Diagnostics). Positivity was determined by the presence of cytoplasmic signals only; nuclear reactivity, either as dot or diffuse, was considered as non-specific staining. Tissue-infiltrated plasma cells expressing high levels of immunoglobulin transcripts acted as internal positive controls.

Human *RAG1* and *RAG2* probes were hybridized using RNAscope 2.5 HD Duplex Reagent Kit (Advanced Cell Diagnostics), adopting an extended 1-hour incubation in Amplifier reagent 5 and 30-minute incubation in Amplifier reagent 6 for *RAG1/RAG2* hybridization. Slide digitalization was achieved using an Aperio CS2 digital scanner (Leica Biosystems) with the ImageScope software (ImageScope version 12.3.2.8013, Leica Biosystems). Quantitative in situ mRNA analysis was performed on whole sections using HALO software (v3.2.1851.229, Indica Labs). A threshold of 20% of cells expressing *RAG1* and/or *RAG2* was used as cut off for positivity.

### In situ Proximity Ligation Assay

Proximity ligation assay (PLA) was performed using NaveniBright HRP kit (Navinci Diagnostics, Uppsala, Sweden) according to manufacturer’s instructions. Polyclonal rabbit anti-human IGM (Leica Biosystems) or rabbit anti-human IGG (Leica Biosystems) were coupled with mouse anti-human CD79A (Leica Biosystems) and mouse anti-human CD79B (developed by B. Falini), respectively. Rabbit anti-human CD79A (Abclonal) was coupled with mouse anti-human CD79B. Stainings with single primary antibodies were used as negative controls. PLA signals were quantified in ten non-overlapping GC areas of human tonsils and five non-overlapping fields of view of IGH^+^ and IGH^UND^ cases at x 200 magnification, using HALO software (v3.2.1851.229; Indica Labs).

### Nucleic acid extraction

DNA was extracted from FFPE tissue specimens using QIAamp DNA FFPE Tissue Kit, according to manufacturer’s instructions (QIAGEN). Amplification of different segments of the *ALB* gene differing in size (300 bp, 484 bp, 600 bp, 800 bp, respectively), served to assess the extent of DNA fragmentation. Concomitant DNA and RNA extraction (AllPrep DNA/RNA Mini Kit; QIAGEN) was achieved from frozen tissue biopsies upon homogenization with stainless steel beads (3–7 mm mean diameter) using TissueLyser II (QIAGEN; 20” at 30 Hz, for 2 cycles(20)), according to manufacturer’s instructions. DNA/RNA concentration and integrity were assessed with Qubit™ dsDNA HS Assay Kit (Thermo Fisher Scientific), NanoDrop™ 2000/2000c Spectrophotometers, and Agilent 2100 Bioanalyzer system.

### Bulk transcriptomics analyses

Polyadenylated RNA was extracted from n=22 lymphoma specimens, including 14 HGBCL-DH-*BCL2*(*-BCL6*) cases. RNA quality and concentration were determined using RNA 6000 NanoChip on 2100 Bioanalyzer (Agilent Technologies). Indexed RNA-seq libraries were prepared from 500 ng of high-quality input RNA (RNA Integrity Number; RIN: >6.5) using a Stranded mRNA ligation kit (Illumina), according to the manufacturer’s instructions. Single-end sequencing (1x75 bp reads, 36 million reads/sample) was used to sequence pooled libraries on a NextSeq 500/550 Illumina sequencer. Raw reads were aligned to the human genome (NCBI build 38, hg38) with STAR v2.7(21) with the parameter --quantMode GeneCounts to count gene reads. Ensemble IDs were changed into gene symbols using BioMArt. The Bioconductor R package DESeq2(22) was used to normalize gene count and to calculate the log2FC between comparisons. The Benjamini-Hochberg method was applied to correct p-values for multiple comparisons. Significantly up/downregulated genes were identified using an adjusted p-value <0.05 and an absolute log2 FC > 0.58. Heatmaps for specific gene signatures were generated with the pheatmap function of the pheatmap R package on DESeq2 normalized counts. IGH isotype choice was inferred from bulk RNA-seq data according to the most expressed *IGH* constant region transcript. FPKM or log2 normalized counts were used for comparisons of gene expression levels, as indicated. Unsupervised interrogation of RNA-seq data from MB2 DE GCB lymphomas for a) DZ/LZ B cell-associated gene expression signatures(23), b) Double-Hit/DZ molecular signature (DHIT/Dzsig (24)), and c) a 370-gene DZ/LZ spatial signature (25), was used to cluster lymphoma samples according to cell-of-origin. After z-score normalization, hierarchical clustering based on the gene signatures listed above was applied to identify clusters reproducing GC heterogeneity. Hierarchical clustering analysis was performed using the Ward.D2 method and considering the Euclidean distances. The rand-index measure was used to quantify the similarity of the two clustering results. Normalized gene expression profiles were subjected to the CIBERSORTx algorithm, to estimate the immune cell subset composition of each lymphoma sample. The LM22 matrix was considered as a reference for immune cell deconvolution. Bulk-mode batch correction (B-mode) was applied for cell fraction estimation (26). The Wilcoxon-Mann-Whitney test was used to compare the CIBERSORTx fractions among conditions.

### Spatial transcriptomics

IGM IHC staining on 4µM thick tissue sections from selected FFPE HGBCL-DH-*BCL2* samples was combined with Regions Of Interest (ROI) definition and segmentation, and *in situ* analysis of 1825 curated cancer- and immune-associated transcripts using the GeoMx Digital Spatial Profiler (DSP) platform (NanoString, Seattle WA), according to manufacturer’s instructions. mRNA binding DNA probes (35-50 nt in size) conjugated with UV photocleavable indexing oligos were hybridized to the tissue as previously reported (27). The UV photocleavable probes were released from each ROI according to custom masks for UV illumination and digitally counted using the NanoString nCounter Analysis System. For nCounter data analysis, digital counts from barcodes corresponding to mRNA probes were normalized to internal spike-in controls. Moreover, a set of six internal housekeeping genes was included in the Human Immuno Oncology panel to control for system variation, including ROI size and cellularity (27,28). Seven negative control probes were adopted to evaluate and filter ROIs with a high degree of non-specific binding, as previously reported (29). Differential expression analyses between IGH^+^ and IGH^UND^ lymphoma ROIs were carried out by applying the moderated t-test using the limma package (30). The Benjamini-Hochberg method was applied to adjust p-values and identify significant differentially expressed genes (adjusted p-value <0.05). The SpatialDecon algorithm (31,32) was used to estimate immunodeconvolution fractions (the safeTME profile matrix was considered as a reference). The Wilcoxon-Mann-Whitney test was used to compare the SpatialDecon fractions among conditions. Fisher’s exact test was applied to assess the significance of the overlap between the gene sets.

### Whole genome sequencing

Whole genome sequencing (WGS) was performed on genomic DNA extracted from FL-HGBCL-DH-*BCL2*(*-BCL6*) metachronous cases. Isolated genomic DNA was quantified with Qubit dsDNA HS Assay Kit (ThermoFisher), and quality was assessed by agarose gel. Library preparation was performed using the KAPA Hyper Prep kit (Roche) per manufacturer’s recommendations. Briefly, gDNA was sheared to 500 bp using Covaris LE220-plus, adapters were ligated, and DNA fragments were amplified with minimal PCR cycles. Library quantity and quality were assessed with Qubit dsDNA HS Assay Kit (ThermoFisher), Tapestation High Sensitivity D1000 Assay (Agilent Technologies), and QuantStudio 5 System (Applied Biosystems). Illumina 8-nt dual indices were used. Equimolar pooling of libraries was performed based on QC values and sequenced on an Illumina NovaSeq S4 (Illumina) with a read length configuration of 150 paired-end for 600M paired-end reads (300M in each direction), at Biodiversa (Rovereto, Italy). Reads were aligned to reference genome (GRCh38) with the Burrows-Wheeler Aligner (BWA-MEM) algorithm (33). Duplicates were removed using the Picard MarkDuplicates tool. To identify *IGHV* rearrangements and translocations targeting IGH/K/L loci, IgCaller (34) was run on each sample using the bam aligned files after duplicates removal and hg38 reference file. A custom script in R was run to identify IGV rearrangements with the same junction in metachronous samples. Genomic coordinates of the breakpoints captured using IgCaller were validated by PCR amplification followed by Sanger sequencing.

### Whole exome sequencing

Genomic DNA was extracted from 5 IGH^+^ and 7 IGH^UND^ lymphoma cases using either QIAamp DNA FFPE Tissue Kit (QIAGEN) (FFPE tissues) or AllPrep DNA/RNA Mini Kit (QIAGEN) (frozen tissues), according to manufacturer’s instructions. Isolated genomic DNA was quantified with Qubit dsDNA HS Assay Kit (ThermoFisher), and quality was assessed by Tapestation genomic DNA Assay (Agilent Technologies).

Library preparation was performed using SureSelectXT Reagent Kit (Agilent Technologies) per manufacturer’s recommendations. Exome capture was performed with the SureSelect Human All

Exon V7 kit (Agilent Technologies). Library quality and quantity were assessed with Qubit dsDNA HS Assay Kit (ThermoFisher), Tapestation High Sensitivity D1000 Assay (Agilent Technologies), and QuantStudio 5 System (Applied Biosystems). Illumina 8-nt dual indices were used. Equimolar pooling of libraries was performed based on QC values and sequenced on an Illumina NovaSeq 6000 with a read length configuration of 150 paired-end for output required, at Biodiversa (Rovereto, Italy). Reads pre-processing was performed using the Genome Analysis Toolkit (GATK4) Best Practices Workflow (35). Reads were aligned to the reference genome (GRCh38) with the BWA-MEM algorithm (33). Removal of duplicates was performed with the MarkDuplicates tool. Base Quality Score Recalibration was achieved by applying the BaseRecalibrator tool with the list of 1000G phase1 high confidence SNPs as known polymorphic sites, followed by the ApplyBQSR tool, which recalibrated the base qualities of the input reads based on the recalibration table produced by the BaseRecalibrator tool. The HaplotypeCaller command of GATK4 was run to call single nucleotide variants and indels. Given the low number of samples, a hard-filtering approach was chosen to filter variants according to quality parameters (“QD <0.2 || MQ <40.0 || FS > 60.0 || SOR > 3.0 || MQRankSum <-12.5 || ReadPosRankSum < −8.0” for SNPs and “QD <0.2 || FS >200.0 || SOR >3.0 || MQRankSum <-12.5||ReadPosRankSum<-20.0” for indels). Functional annotation of variants was performed with the FUNCtional annOTATOR (Funcotator) tool of GATK4 using the germline hg38 database (v1.4.20180829g) for gene annotation, variant classification, and DBSNP frequency. The SnpSift(36) tool was applied for annotation of SNPs present in the Single Nucleotide Polymorphism database using as a reference file the All_20180418.vcf file downloaded from ftp://ftp.ncbi.nih.gov/snp/organisms/human_9606/VCF/.

The Annovar(37) software was utilized to infer the pathogenic score predictions using the ljb26 database. Variant filtering was performed in R, after the merging of Funcotator, SnpSift, and Annovar outputs. Variants with read depth (DP) <10 in all the samples or with allele frequency (AF) <0.05 in all samples were removed from the analysis. Variants annotated in dbSNPs with a minor allele frequency >5% in more than one population also were filtered out.

### *IGV* gene rearrangement analysis

*IGHV* rearrangements were PCR amplified from genomic DNA extracted from FFPE tissue sections or frozen biopsies of selected lymphoma cases, followed by deep-sequencing on the Illumina MiSeq platform, using a modified version of the two-step PCR-based BIOMED-2 protocol(38). As first-step PCR amplification, a set of forward primers annealing to human *IGHV* genes was coupled with reverse primers annealing to human *IGHJ* segments. All primers were adapted for compatibility with the Illumina Indexing set (Nextera XT Index Kit - 96 indexes, Illumina) (Primer sequences are listed in Supplementary table S7). First-round PCR products were purified using AMPure XP Beads (Beckman Coulter) before the second-round indexing PCR and library preparation steps. *IGHV* rearrangements were also profiled using the LymphoTrack Dx *IGHV* Leader Somatic Hypermutation Assay for Illumina MiSeq (Invivoscribe), according to manufacturer’s instructions. The concentration and quality of libraries were determined by Qubit dsDNA HS Assay Kit (ThermoFisher) and the 2100 Bioanalyzer platform (Agilent), respectively. Libraries were sequenced using MiSeq Reagent Kit v3, 2.300bp (Illumina) or MiSeq Reagent Kit v2, 2.300bp (Illumina) (2 × 250 paired-end reads), on the MiSeq Illumina platform. Illumina PhiX spike-in was added to low-complexity libraries (Illumina). *IGHV* sequencing reads were analyzed using the Immcantation framework (https://immcantation.readthedocs.io/en/stable/). Assembly of the paired reads, quality filter, and primer mask were performed using the pRESTO toolkit(39). Each set of paired ends reads was first assembled into a full-length *IGV* sequence using the subcommand AssemblePairs.py of pRESTO. The FilterSeq.py command was then used to remove low-quality reads with a Phred quality score of less than 20, and the MaskPrimers.py was applied to identify and remove PCR primers annealing to V- and C-regions for both reads. The CollapseSeq.py command was used with the “-n 20 --inner --uf CPRIMER --cf VPRIMER --act set” parameters to identify unique sequences. Following duplicate removal, unique sequences with at least two representative reads were selected by using the command SplitSeq.py on the count field (-f DUPCOUNT) and a threshold of 2 (--num 2). Reads were subsequently aligned to the IMGT human IG *V*, *D,* and *J* reference sequences with IgBlast using the igblastn command line. The ParseDb.py command of Change-O(40) was run with the option “select” to subset heavy chain reads. Clonality was determined with the DefineClones.py command of Change-O that was run with the following parameters: “--act set --model ham --norm len --gf d_call --mode allele -- maxmiss 6 --link average” and using an appropriate threshold determined with the distToNearest function of the SHazaM package, that calculates the distance between each sequence in the data and its nearest-neighbor. The latter function was also called with model=“ham” to use a single nucleotide Hamming distance matrix with gaps assigned zero distance and normalize=“len” to normalize the distance by the length of the sequence group. The union of ambiguous gene assignments was used to group all sequences with any overlapping gene calls.

### IGV tree-based mutational profiling, lineage tree and selection analyses

*IGV* lineage tree and selection analyses were performed on selected IGH^+^ and IGH^UND^ cases. High-throughput sequencing of *IGHV* reads was pre-processed using pRESTO version 0.5.13 (39). The pre-processing included assembly of paired ends, quality filtering by trimming low-quality edges, filtering out reads with an average Phred score lower than 25, and masking bases with Phred scores lower than 20. Sequences with more than 10 masked or missing bases were discarded. FWR1-region and J-region primers (38) were masked (replaced by N’s) to preserve gene length. Next, identical sequences were collapsed, and only sequences with two copies or more were selected for analysis; the selected sequences were processed using Change-O version 0.4.6(40) and in-house custom scripts. The processing included sequence annotation with IMGT/GENE-DB (Giudicelli et al. (41); germline segment sequences were downloaded from IMGT on July 1, 2021), clonal assignment, and assessments of sampling depth and clonal size distributions. Putative germline sequences for each clone were created based on the same IMGT/GENE-DB database and clonal consensus in junction regions. The largest (dominant) clone identified in each repertoire was assumed to be the tumoral clone. Samples with polyclonal profiles were omitted from the analysis. Clones containing more than two unique sequences from the dominant, non-dominant, or healthy control groups were analyzed using IgTree© (42) for lineage tree construction, and lineage tree-based analyses were performed using IgTreeZ (43). To exclude the pre-transformation mutational history of each lymphoma clone from the analysis, trunks were removed from the IGH^UND^ and IGH^+^ trees, and the first split node of each “trunkless” tree was assigned to be the new root node. Trees that originally had no trunks were removed from the trunkless analysis to avoid bias due to trees not having enough information regarding their diversification history.

Visual representations of lineage tree shapes were created using the graph description language DOT, as implemented in Graphviz (44). Node (sequence) names were omitted for better tree visualization. Selection analysis was performed using the program BASELINe, as implemented in the SHazaM package (40,45). The BASELINe algorithm was used to a) calculate the expected mutation frequency, b) estimate the selection strength for each clonal lineage tree, and c) compare the selection scores of the repertoires in each group of clones. *IGHV* repertoires from peripheral blood mononuclear cells isolated from healthy donors (46) were used as controls.

Comparisons between lymphoma lineage tree characteristics with those of healthy repertoires, which included more than 50,000 trees, were performed based on patient average, to overcome the bias of the control dataset being so much larger. Comparisons against clones from Reactive LN, which included 599 trees from two patients, were done based on trees, as two samples are insufficient for statistical inference. For each comparison, the assumptions of normal data distribution and variance homogeneity were tested using the Shapiro test and the Levene test, respectively. If the data were normally distributed and had homogenous variances, Student’s t-test was used. Otherwise, the non-parametric Mann–Whitney U-test was used. To correct for multiple comparisons, Benjamini and Hochberg’s False detection rate (FDR) method was applied.

### Quantification of *IGHM* copy number

Genomic DNA was extracted using AllPrep DNA/RNA Mini Kit (QIAGEN) according to manufacturer’s protocol. Human *IGHM* constant region gene sequence was obtained from GenBank (https://www.ncbi.nlm.nih.gov/genbank). Primers annealing to *IGHM* were designed using Primer3plus tool and are listed in Supplementary table S7. Quantitative genomic real-time PCR (gRT-PCR) reaction was performed using TB Green Premix Ex Taq TM II (Tli RNaseH Plus; Takara) following standard procedures, on the Light Cycler 480 II (Roche Diagnostics). Gene copy number was determined for HGBCL-DH-*BCL2(-BCL6)* samples after normalization for DNA input, testing a segment of *CD3G* as a reference, using the comparative Ct method (2^-ΔΔCt method). Tonsillar IGM^+^ human B cells from healthy donors were used as a reference.

### Quantification of *RAG1* and *RAG2* transcripts

Total RNA was extracted from COH-THL1, COH-DHL1, WILL-2, NAMALWA, NU-DHL-1, NU-DUL-1, SU-DHL-8, DoGKIT, SU-DHL-4, DOGUM, 697, and 293T human B lymphoma cell lines, and λ-MYC;B1-8f murine lymphoma lines, using RNeasy Mini extraction kit (QIAGEN), following manufacturer’s instructions. Total RNA was reverse transcribed into complementary DNA (cDNA) using Superscript III Reverse Transcriptase (ThermoFisher Scientific) according to manufacturer’s instructions. Quantitative real-time PCR (qRT-PCR) reactions were performed using TB Green Premix Ex Taq TM II (Tli RNaseH Plus; Takara) on a Light Cycler 480 II (Roche Diagnostics). *GAPDH* gene served as control(47). Human and murine *RAG1*, *RAG2*-specific and GAPDH primers are listed in Supplementary table S7.

### 5’ Rapid Amplification of cDNA Ends (RACE)

5’ Rapid Amplification of cDNA Ends (5’RACE) protocol was adapted from Bernat et al.(48). Briefly, total RNA was extracted using AllPrep DNA/RNA Mini Kit (QIAGEN), and first-strand cDNA synthesis reaction was performed on 200 ng of total RNA with 10 µM oligo dT and Superscript II Reverse Transcriptase (ThermoFisher Scientific) at 42^◦^C for 1 h. Template switching was performed by adding to the reaction mix a template switch oligonucleotide (Read1_TS Supplementary table S7) composed of a 5 nucleotide poly-guanosine (poly-G) stretch, a 12 nucleotide unique molecular identified (UMI) and an Illumina’s universal amplification sequence (Read1). The template switch reaction was carried out at 42^◦^C for 1 h. cDNA was purified using the Wizard SV Gel and PCR clean-up system (Promega), according to manufacturer’s instructions. 2X KAPA HiFi Hot Start Ready Mix (Roche) was used to amplify the purified cDNA template through a 2-step semi-nested PCR using forward Read1-specific primers (Read1-U) and 2 sets of class-specific *IGH* and *lGK/L* constant region gene-specific reverse primers (with second-round primers including Illumina Read2 sequence). Primer sequences are listed in Supplementary table S7. A purification step was conducted after each PCR round using AMPure XP Bead-Based Reagent (Beckman Coulter Life Sciences), according to manufacturer’s instructions, using a 0.65X beads-to-sample ratio. 10 ng of purified PCR products were indexed using 2 x KAPA HiFi Hot Start Ready Mix, with forward (P5_R1) and reverse (P7_R2) indexing primers, in a 13-cycle index PCR reaction. Indexed libraries were purified using AMPure XP Beads, quality controlled using Agilent 2100 Bioanalyzer system, and sequenced (2 × 250 bp read length) on a MiSeq Illumina sequencer, using MiSeq Reagent Kit v2, 2.300bp (Illumina).

Fastq files were merged with usearch with standard parameters (v.11.0.667, -fastq_minmergelen 15, -fastq_pctid 90). 3’ primers were removed with cutadapt and sequences not containing 3’ primers were discarded (v.3.4, --overlap 20, -e 0.05, --discard-untrimmed). Primers-removed sequences were collapsed with an in-house Python package; sequences observed only once were discarded. Fasta sequences were aligned with Igblast (v.1.21.0) against the IMGT database. Further analyses were performed with the Immcantation change-o package (v.1.3.0 (40)). To avoid wrong V-gene assignment during clonotypes identification due to an intrinsic heterogeneity in RACE’s amplicon size, a Python-based clonotypes assigner was used to integrate differences in V-gene assignment, weighting them through hamming distance calculation. Sequences with identical CDR3 length and D-J-gene assignment were grouped to perform CDR3 clustering analysis. CDR3 nucleotide distances were calculated among grouped sequences, and precomputed V-gene weights were added depending on the IgBlast V-gene assignment. Agglomerative clustering was performed to cluster the same clonotype sequences (sklearn package AgglomerativeClustering, metric=’precomputed’, n_clusters=None, linkage=’average’, distance_threshold =cutoff). A dynamic distance cutoff based on CDR3 lenght (cutoff = 0.15 if cdr3 length <=30,cutoff = 0.09 if cdr3 length <=51, cutoff = 0.08 if cdr3 length > 51) was applied.

### Analysis of *IGK KDE* rearrangements by genomic PCR

Genomic DNA was extracted from HeLa, PBMC, COH-DHL1, COH-THL1 IGK^+^, COH-THL1 IGL^+^, and COH-THL1 IGH^UND^ cell lines using AllPrep DNA/RNA Mini Kit (QIAGEN), following manufacturer’s instructions. *IGKV* family-specific forward primers, together with an oligonucleotide annealing to the genomic region between *IGKJ* and *IGKC* segments, were combined with a *KDE* reverse primer to amplify *KDE* rearrangements(49). PCR reactions were carried out with GoTaq Flexy polymerase kit (Promega), using the SimpliAmp Thermal Cycler (Thermo-Applied Biosystem). Primers amplifying a segment of the I*GKC* gene were designed to assess bi-allelic *KDE* rearrangements by genomic PCR. PCR amplification of a segment of the *RPLPO* gene served to control DNA input (50). Oligonucleotide sequences are listed in Supplementary table S7.

### Treatment of lymphoma cell lines with Polatuzumab-Vedotin

HT (IG^UND^), RAMOS (IGM^+^), DoGKIT (IG^UND^), COH-THL1 (IGM^+^), COH-THL1 (IGH^UND^), and COH-DHL1 (IGH^UND^) human lymphoma lines were treated with Polatuzumab (P)-Vedotin (V) or Polatuzumab (P) (MedChemExpress). PV was dissolved in PBS, and stored in aliquots at −80°C. Cells were seeded in a 6-well plate at a density of 5 x 10^5 cells/ml and treated with PV or P, at a concentration of 1.25 µg/ml. All experiments were performed at least in triplicate. Vehicle-treated controls were included in each experiment. Cell viability was assessed 72 h after treatment using LIVE/DEAD™ Fixable Aqua Dead Cell Stain Kit (ThermoFisher Scientific).

### Animal models

BCR-positive and BCR-negative λ-MYC;B1-8f lymphoma cells were isolated from two independent primary tumors (#2646 and #2567) from 12- to 20-week-old male and female λ-MYC;B1-8f mice on a C57BL/6J x BALB/c mixed genetic background, as previously described(5). 8-12-week-old male immunocompetent syngeneic CB6F1/J mice were used as recipients for tumor transfer experiments. All animals were housed following institutional and national guidelines and regulations. Animal procedures and studies were performed in compliance with Italian national and EU directives (2010/63/EU) for animal research with protocols approved by the local ethical committee and the Italian Ministry of Health (# 561/2021-PR).

### In vivo lymphoma studies

BCR-negative murine λ-MYC;B1-8f lymphoma cells were in vitro-derived from their BCR^+^ counterparts by TAT-Cre transduction starting from two independent primary tumors (#2646 and #2567), as previously described (5). Briefly, BCR^+^ lymphoma B cells were transduced with TAT-Cre (37.5 μg/mL) protein in serum-free media (Hyclone) at 5×10^6^ cells/mL for 45 min, at 37 °C. Transduced cells were washed three times and cultured using complete B cell medium (high-glucose DMEM supplemented with 10% heat-inactivated FBS, 0.1 mM nonessential amino acids, 1 mM sodium pyruvate, 50 μM β-mercaptoethanol and 2 mM L-glutamine). BCR^−^ lymphoma cells were isolated by negative selection through magnetic-activated cell sorting (MACS), using biotin-labeled anti-mouse IgM (cl.# R33.24.12), followed by anti-biotin microbeads (Miltenyi Biotech), or by cell sorting using FACS Aria Instrument (BD Pharmingen, USA) upon staining with a monovalent fluorescently labeled anti-mouse IgM (Fab, Jackson Immunoresearch). 1.5 million purified BCR^+^ or BCR^-^ lymphoma cells were transferred intravenously into 8-12-week-old male immunocompetent syngeneic CB6F1/J recipients. Transplanted animals were sacrificed 14 (BCR^+^) or 21 (BCR^-^) days post-transfer. Flow cytometry analysis was used to assess the infiltration of IgM^+^ or IgM^-^ lymphoma B cells (FSC^hi^, CD19^+^, MYC^+^) in primary and secondary lymphoid organs. FSC^hi^/CD19^+^ BCR^+^ or BCR-less lymphomas were FACS-purified using a monovalent (Fab) anti-IgM antibody to prevent antigen receptor crosslinking. DNA from BCR^+^ and BCR^-^ ex-vivo purified cells was extracted using genomic DNeasy tissue kit (QIAGEN). Genomic PCR amplification of the *B1-8f* allele (present in BCR^+^ while lacking in BCR^-^ derivatives) was used to BCR genotype lymphomas cells. VDJ profiling in *B1-8* knock-out *(Δ)* lymphoma cells spontaneously restoring IgM expression was performed using primers previously described (51). Briefly, DNA amplification was carried out in two rounds of nested PCR approach using GoTaq G2 DNA Polymerase (Promega). Two degenerate forward primers annealing to framework region 1 of *IGHV* genes and a degenerate reverse primer annealing to *IGHJ* segments were used for the first-round amplification (52). In the second PCR reaction, 1 μl of the first-round amplicon was reamplified with forward primers annealing to *IGHV* family-specific primers (*J558, GAM3, 36-60, S107, 71835, X-24, Q52*), in combination with an *IGHJ4*-specific reverse primer. PCR amplicons were cloned into pGEM-Teasy vector, transformed in DH5α bacteria, and plasmid DNA subjected to Sanger sequencing. *IGHV* analysis was performed using IGBLAST.

### Quantification and statistical analysis

No statistical methods were used to predetermine the sample size. GraphPad Prism software v.10.1.1 was used to graphically represent data and perform statistical analyses. Unpaired Student’s t-test, non-parametric Mann-Whitney test, Fisher’s Test, and Chi-square test were used to analyze data, as indicated. The results section, figures, and figure legends report information on sample size, mean and median values, and the statistical test used. For all analyses, p values <0.05 were considered significant.

## Resource availability

Further information and requests for resources, reagents or materials should be directed to and will be fulfilled by the corresponding author.

## Data availability

Data generated in this study will be available following article publication and/or upon request directed to the corresponding author.

