## Supplementary Figures for "B cell receptor silencing reveals the origin of high-grade B cell lymphomas with *MYC* and *BCL2* rearrangements"

### Figure S1

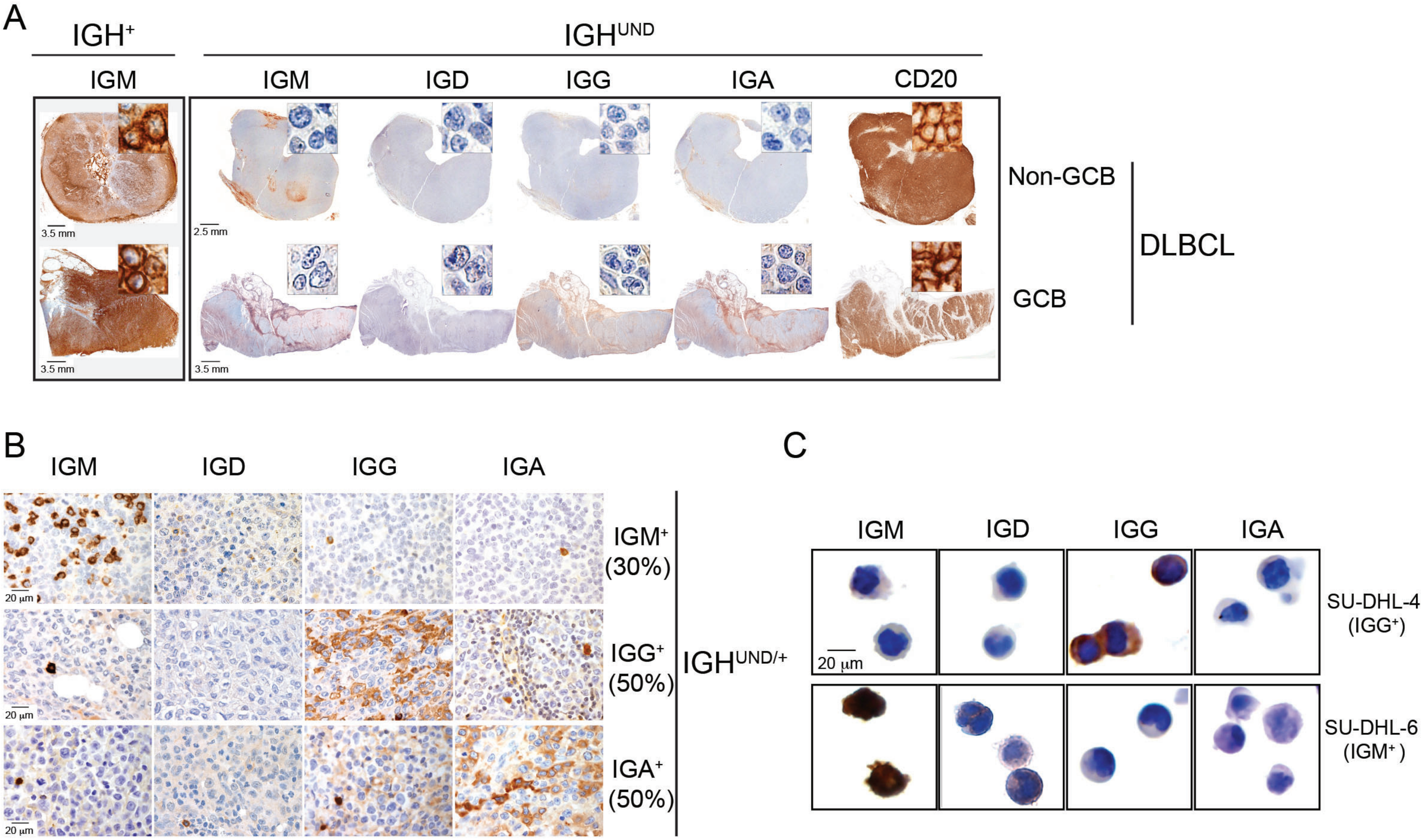
A
IGH+
IGHUND
IGM
IGM
IGD
IGG
IGA
CD20
3.5 mm
2.5 mm
3.5 mm
3.5 mm
Non-GCB
GCB
DLBCL
B
IGM
IGD
IGG
IGA
20 µm
20 µm
20 µm
IGM+ (30%)
IGG+ (50%)
IGA+ (50%)
IGHUND/+
C
IGM
IGD
IGG
IGA
20 µm
SU-DHL-4 (IGG+)
SU-DHL-6 (IGM+)

### Figure S2

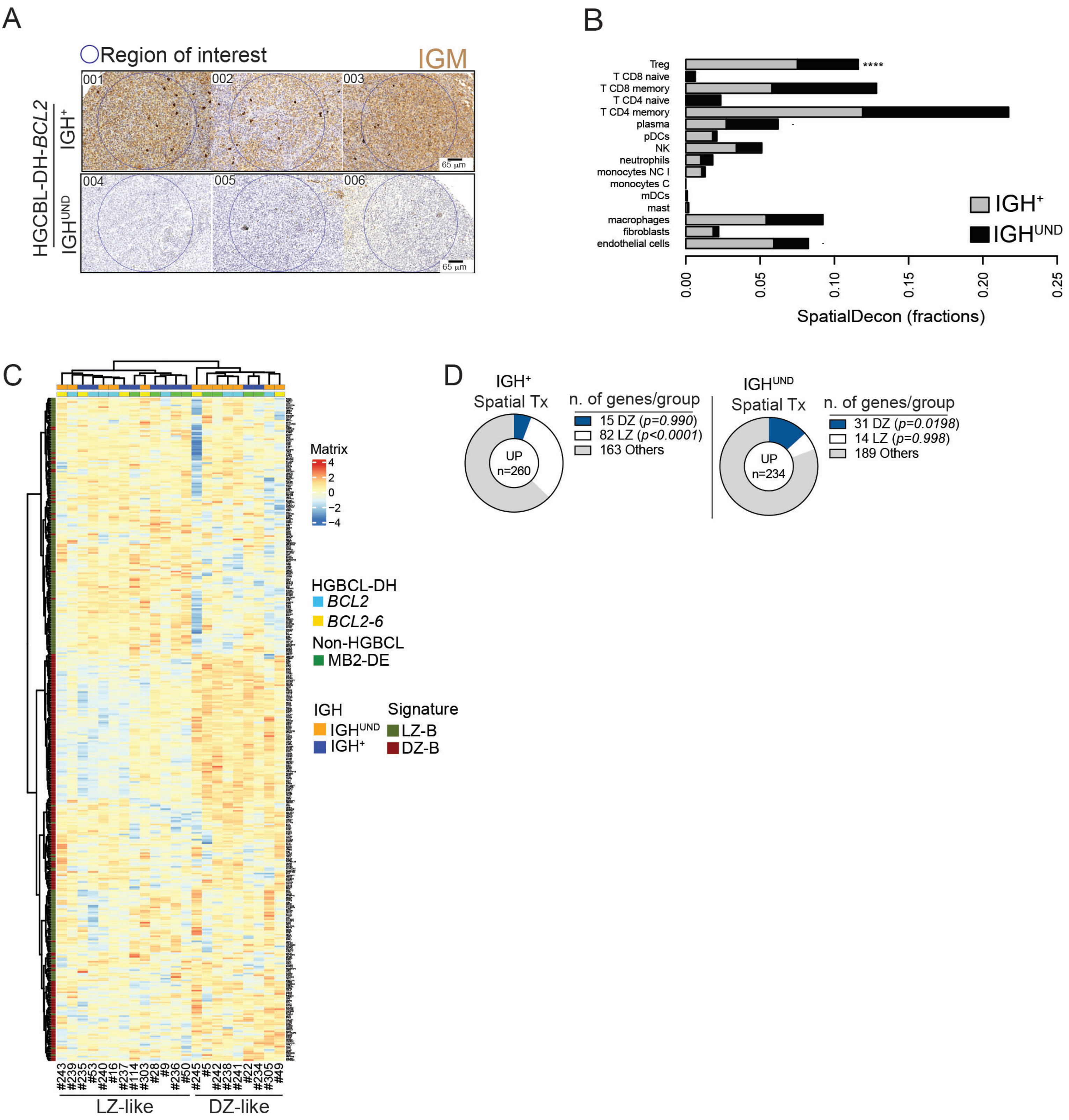

A
Region of interest
IGM
001
002
003
004
005
006
65 µm
HGCBL-DH-BCL2
IGH+
IGHUND
B
Treg
T CD8 naive
T CD8 memory
T CD4 naive
T CD4 memory
plasma
pDCs
NK
neutrophils
monocytes NC I
monocytes C
mDCs
mast
macrophages
fibroblasts
endothelial cells
****
IGH+
IGHUND
0.00
0.05
0.10
0.15
0.20
0.25
SpatialDecon (fractions)
C
Matrix
4
2
0
−2
−4
HGBCL-DH
BCL2
BCL2-6
Non-HGBCL
MB2-DE
IGH
IGHUND
IGH+
Signature
LZ-B
DZ-B
#243
#239
#235
#53
#240
#16
#237
#114
#303
#28
#9
#236
#50
#245
#5
#242
#238
#241
#22
#234
#305
#49
LZ-like
DZ-like
D
IGH+
Spatial Tx
UP
n=260
n. of genes/group
15 DZ (p=0.990)
82 LZ (p<0.0001)
163 Others
IGHUND
Spatial Tx
UP
n=234
n. of genes/group
31 DZ (p=0.0198)
14 LZ (p=0.998)
189 Others

### Figure S3

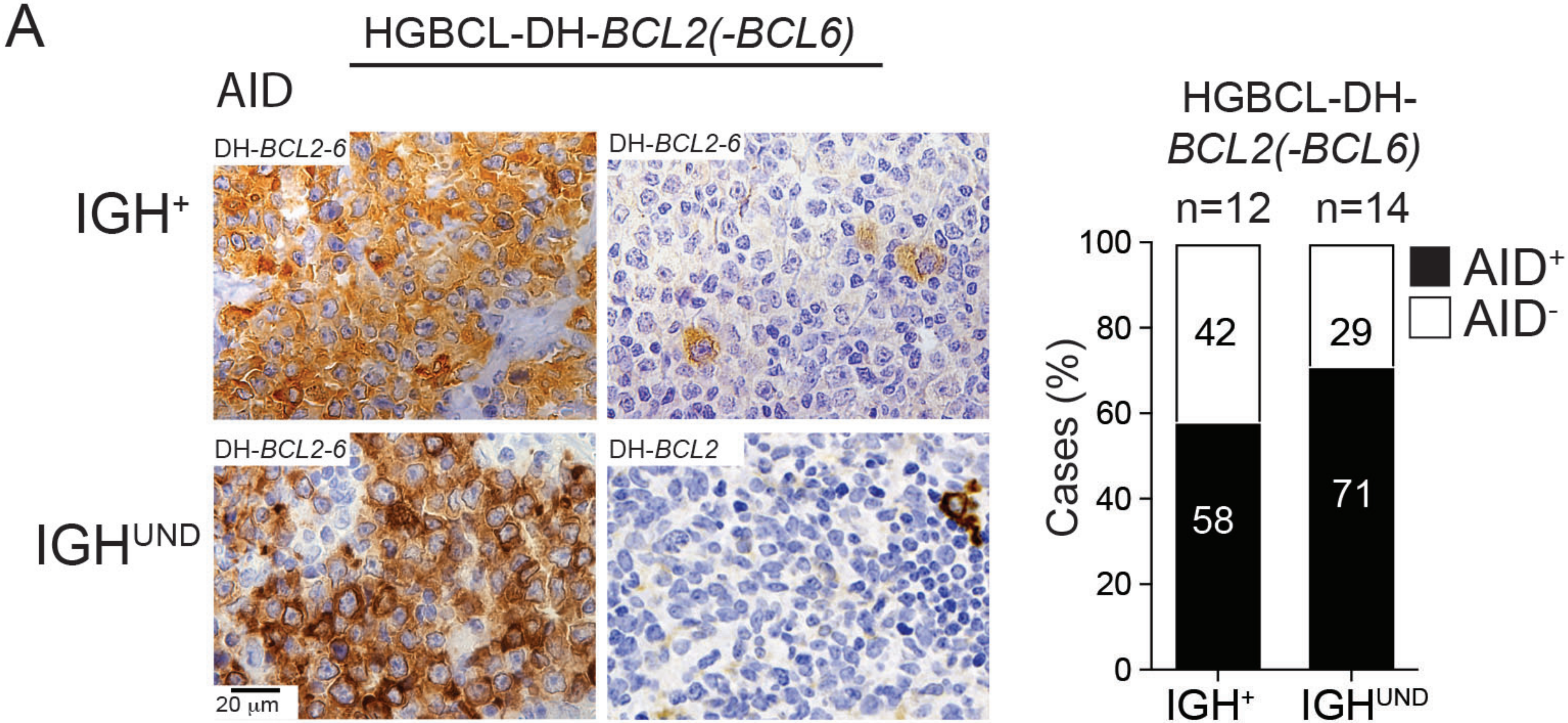
A
HGBCL-DH-BCL2(-BCL6)
AID
IGH+
IGHUND
DH-BCL2-6
DH-BCL2-6
DH-BCL2-6
DH-BCL2
20 µm
HGBCL-DH-BCL2(-BCL6)
n=12
n=14
AID+
AID-
Cases (%)
100
80
60
40
20
0
42
29
58
71
IGH+
IGHUND

### Figure S4

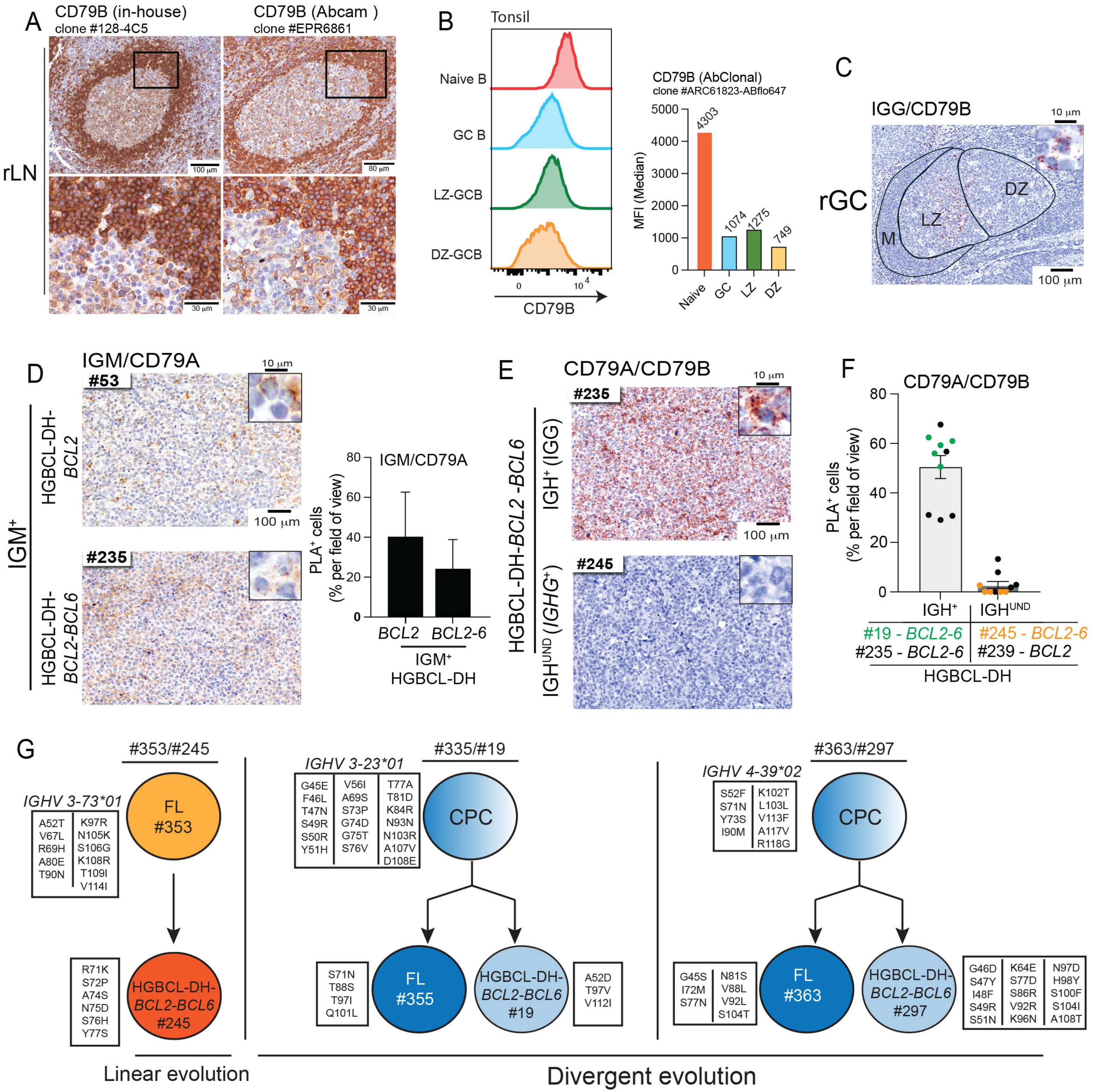

A
CD79B (in-house)
clone #128-4C5
CD79B (Abcam )
clone #EPR6861
rLN
B
Tonsil
Naive B
GC B
LZ-GCB
DZ-GCB
CD79B
CD79B (AbClonal)
clone #ARC61823-ABflo647
MFI (Median)
4303
1074
1275
749
Naive
GC
LZ
DZ
C
IGG/CD79B
rGC
LZ
DZ
M
D
IGM/CD79A
#53
#235
IGM+
HGBCL-DH-BCL2
HGBCL-DH-BCL2-BCL6
PLA+ cells (% per field of view)
BCL2
BCL2-6
IGM+ HGBCL-DH
E
CD79A/CD79B
#235
#245
HGBCL-DH-BCL2-BCL6
IGH+ (IGG)
IGHUND (IGHG+)
F
CD79A/CD79B
IGH+
IGHUND
#19 - BCL2-6
#235 - BCL2-6
#245 - BCL2-6
#239 - BCL2
HGBCL-DH
G
#353/#245
IGHV 3-73*01
A52T V67L R69H A80E T90N
K97R N105K S106G K108R T109I V114I
FL #353
R71K S72P A74S N75D S76H Y77S
HGBCL-DH-BCL2-BCL6 #245
Linear evolution
#335/#19
IGHV 3-23*01
G45E F46L T47N S49R S50R Y51H
V56I A69S S73P G74D G75T S76V
T77A T81D K84R N93N N103R A107V D108E
CPC
S71N T88S T97I Q101L
FL #355
HGBCL-DH-BCL2-BCL6 #19
A52D T97V V112I
#363/#297
IGHV 4-39*02
S52F S71N Y73S I90M
K102T L103L V113F A117V R118G
CPC
G45S I72M S77N
N81S V88L V92L S104T
FL #363
HGBCL-DH-BCL2-BCL6 #297
G46D S47Y I48F S49R S51N
K64E S77D S86R V92R K96N
N97D H98Y S100F S104I A108T
Divergent evolution

### Figure S5

## A

#### *IGH* Break-Apart (BAP) FISH pattern (not in scale)

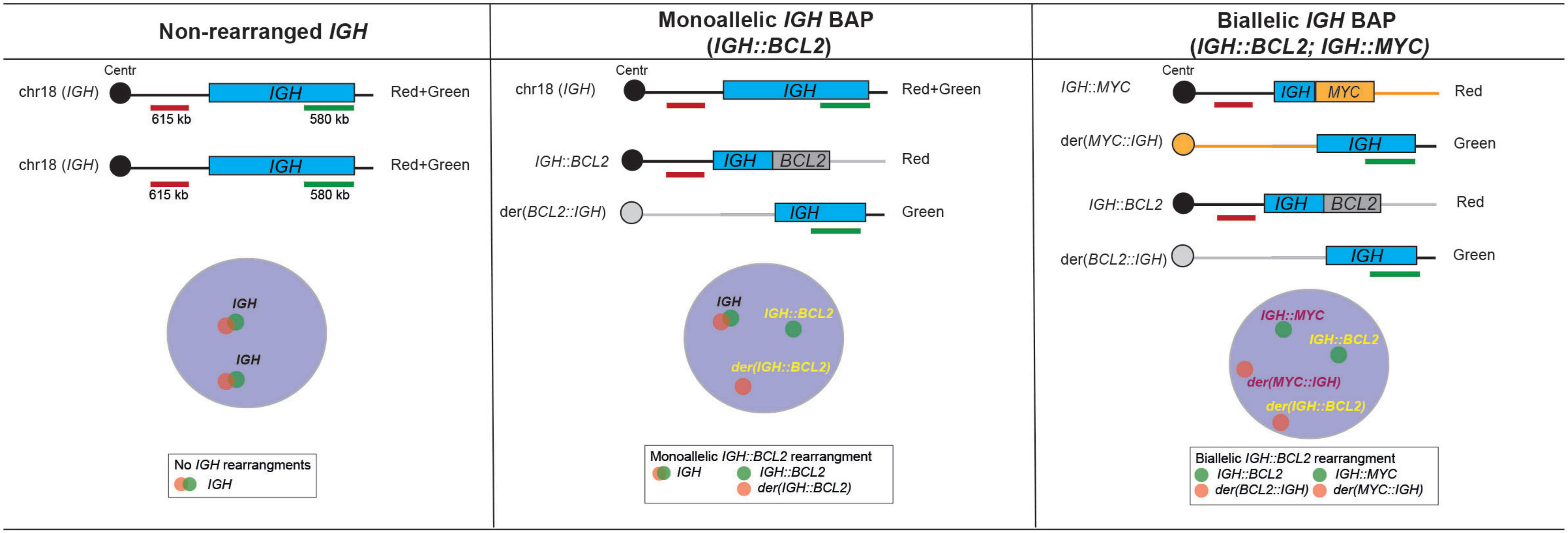


## B

#### *IGH::BCL2* dual color dual fusion pattern (not in scale)

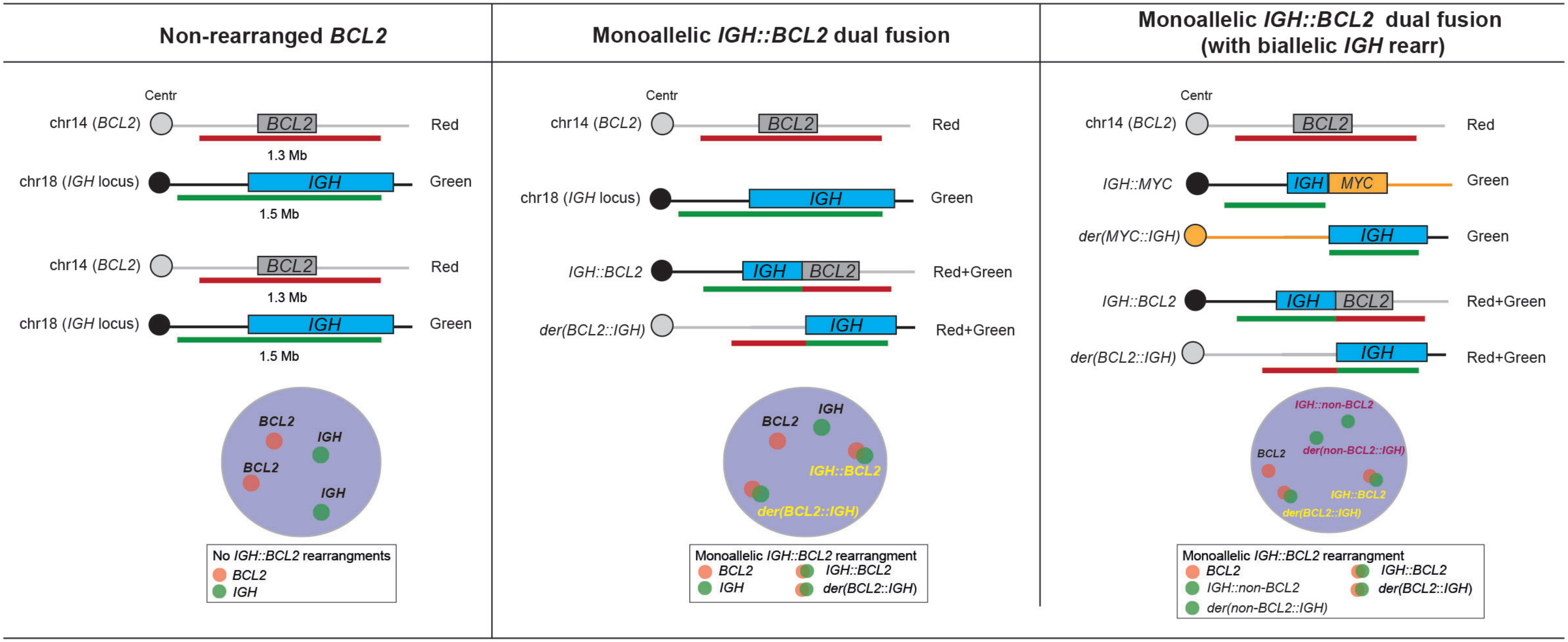

Figure S6

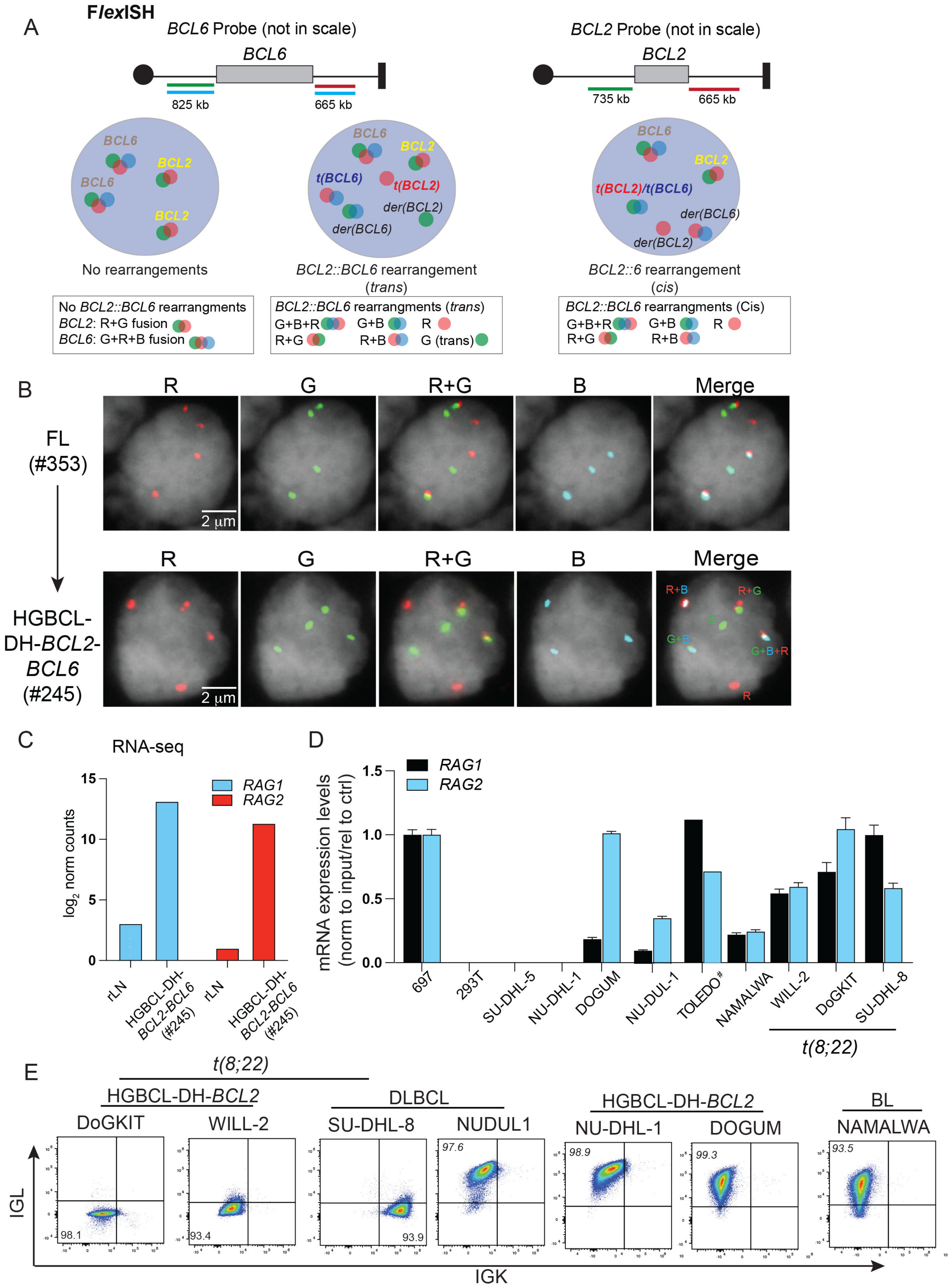

A
FlexISH
BCL6 Probe (not in scale)
BCL6
825 kb
665 kb
BCL2 Probe (not in scale)
BCL2
735 kb
665 kb
No rearrangements
BCL2::BCL6 rearrangement (trans)
BCL2::6 rearrangement (cis)
No BCL2::BCL6 rearrangments
BCL2: R+G fusion
BCL6: G+R+B fusion
BCL2::BCL6 rearrangments (trans)
G+B+R G+B R
R+G R+B G (trans)
BCL2::BCL6 rearrangments (Cis)
G+B+R G+B R
R+G R+B
B
R
G
R+G
B
Merge
FL (#353)
HGBCL-DH-BCL2-BCL6 (#245)
2 µm
C
RNA-seq
log2 norm counts
RAG1
RAG2
rLN
HGBCL-DH-BCL2-BCL6 (#245)
D
mRNA expression levels (norm to input/rel to ctrl)
RAG1
RAG2
697
293T
SU-DHL-5
NU-DHL-1
DOGUM
NU-DUL-1
TOLEDO#
NAMALWA
WILL-2
DoGKIT
SU-DHL-8
t(8;22)
E
t(8;22)
HGBCL-DH-BCL2
DoGKIT
WILL-2
DLBCL
SU-DHL-8
NUDUL1
HGBCL-DH-BCL2
NU-DHL-1
DOGUM
BL
NAMALWA
IGL
IGK

### Figure S7

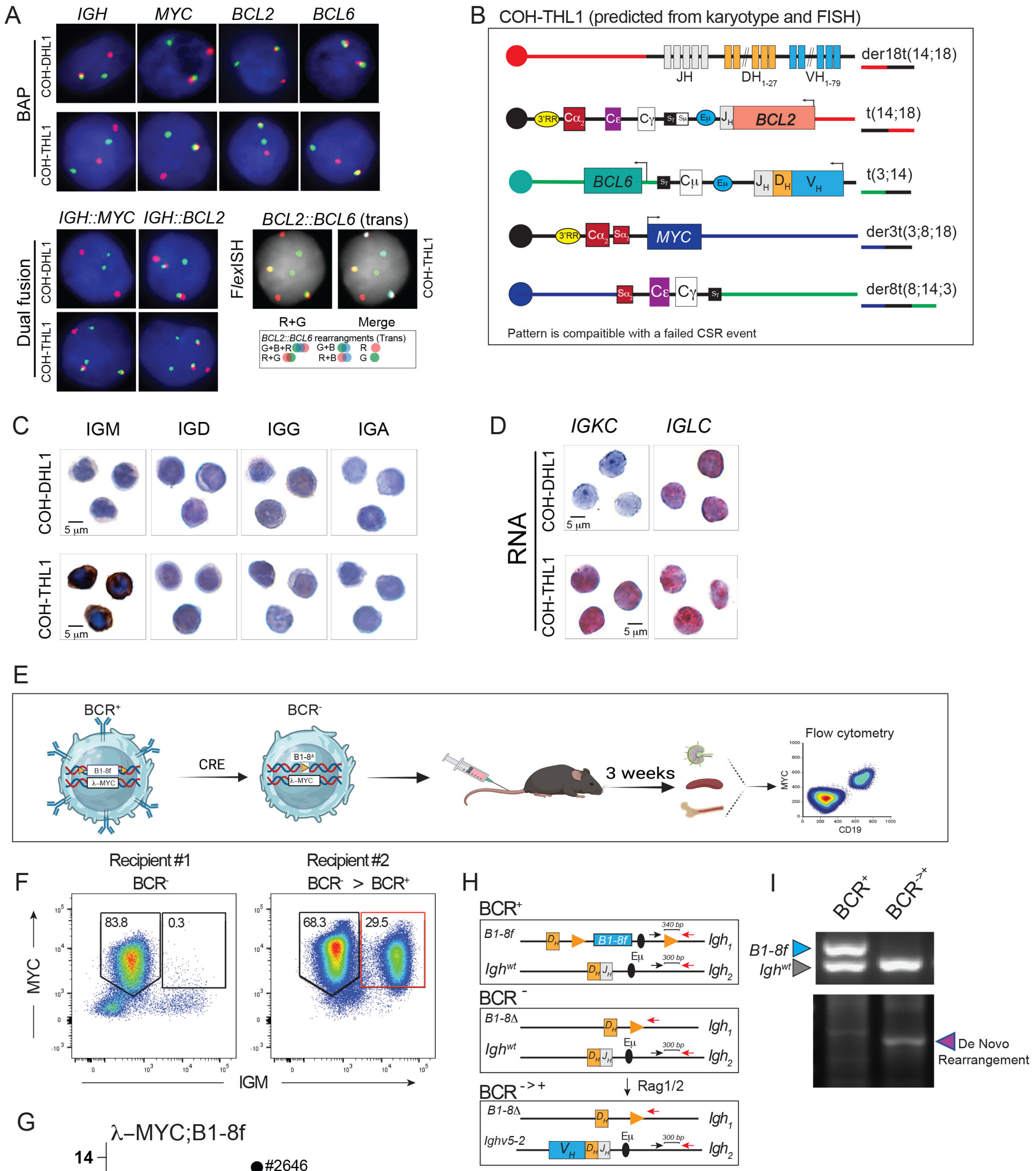


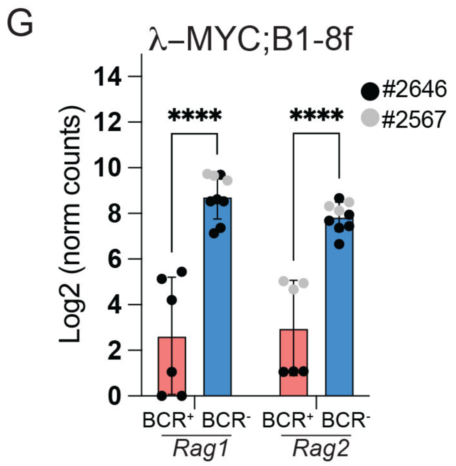


J

VDJ profiling

| Sample | Top V gene match | Top D gene match | Top J gene match | V-J frame | N of sequence | Productive |
|---|---|---|---|---|---|---|
| BCR⁻→BCR⁺ | IGHV5-2*01 | N/A | IGHJ3*01 | In-frame | 12 | Yes |
| | IGHV5-17*01 | IGHD2-4*01,IGHD2-9*02 | IGHJ3*01 | In-frame | 5 | Yes |
