## Supplementary Figure legends for "B cell receptor silencing reveals the origin of high-grade B cell lymphomas with *MYC* and *BCL2* rearrangements"

**Supplemental information titles and legends**

**Supplemental figures**

Supplementary Figure S1. Class-specific IGH expression in DLBCL real-world cases and cell lines.

Supplementary Figure S2. Spatial transcriptomics of $IGH^{+}$ and $IGH^{UND}$ HGBCL-DH-*BCL2*.

Supplementary Figure S3. AID expression in $IGH^{+}$ and $IGH^{UND}$ HGBCL-DH-*BCL2(-BCL6)*.

Supplementary Figure S4. Protein levels of CD79B, and formation of complexes with IGH and CD79 in reactive lymphoid tissues and HGBCL-DH-*BCL2(-BCL6)*.

Supplementary Figure S5. Interpretation of *IGH* BAP and *IGH::BCL2* FISH signals.

Supplementary Figure S6. RAG1/2 expression in $IGH^{UND}$ HGBCL-DH-*BCL2* cell line models.

Supplementary Figure S7. Inducible *IgH* extinction in λ-MYC mouse B cell lymphomas triggers RAG1/2-dependent BCR revision in vivo.

**Supplementary tables**

Supplementary Table S1. DLBCL cases included in the study. Three tables (S1A-C) are included.

Supplementary Table S2. Bulk and spatial gene expression data. Eight tables (S2A-H) are included.

Supplementary Table S3. List of SNVs in HGBCL-DH-*BCL2(-BCL6)*, *IGHV* rearrangements and AID expression status in HGBCL-DH-*BCL2(-BCL6)*. Three tables (S3A-C) are included.

Supplementary Table S4. *IGH* transcript and BCR complex determination in HGBCL-DH-*BCL2(-BCL6)***.** Seven tables (S4A-G) are included.

Supplementary Table S5. *IGH, MYC* and *BCL2* loci in HGBCL-DH-*BCL2(-BCL6)*.

Supplementary Table S6. *RAG1/2* levels and *IGK/LV* rearrangements in HGBCL-DH-*BCL2(-BCL6)* primary specimens and cell lines. Seven tables (S6A-G) are included.

Supplementary Table S7. Key Resources Table. Four tables are included.

**Supplementary data**

**Supplementary Figure S1. Class-specific IGH expression in DLBCL cases and cell lines**

(A) Whole section tracking of class-specific IGH expression on FFPE sections of representative $IGH^{+}$ and $IGH^{UND}$ DLBCL cases, grouped according to Hans classification, assessed by IHC. CD20 staining acted as an internal control. Insets reveal IGH expression in individual cells of representative neoplastic areas.

(B) Representative DLBCL $IGH^{UND/+}$ mixed cases assessed for class-specific IGH by IHC analysis. Frequency of $IGH^{+}$ cells was estimated scoring >100 cells in at least n=3 fields of view.

(D) IHC analyses for the indicated IGH chains of FFPE cytoblocks of SU-DHL4 ($IGG^{+}$) and SU-DHL-6 ($IGM^{+}$) lymphoma cells.

**Supplementary Figure S2. Spatial transcriptomics of $IGH^{+}$ and $IGH^{UND}$ HGBCL-DH-*BCL2***

(A) IGM IHC analysis of regions of interest (circles) of $IGH^{+}$ and $IGH^{UND}$ HGBCL-DH-*BCL2* FFPE specimens, profiled by GeoMx digital spatial profiler, photographed at high magnification.

(B) Cell-type deconvolution according to SpatialDecon, comparing $IGH^{+}$ (grey bars) to $IGH^{UND}$ (black bars) HGBCL-DH-*BCL2* ROI (**** $p < 2e\text{-}04$).

(C) Unsupervised clustering of $IGH^{+}$ and $IGH^{UND}$ MB2 DE DLBCL cases for GC DZ/LZ B cell signatures. Heatmap identifies genes preferentially expressed in DZ (red) or LZ (green) B cells.

(D) Intersection between transcripts upregulated in ROI of $IGH^{+}$ or $IGH^{UND}$ HGBCL-DH-*BCL2* cases and spatially-resolved GC DZ/LZ gene signatures. Significant ($p < 0.0001$) enrichment for GC LZ-associated transcripts was observed for genes preferentially expressed in $IGH^{+}$ HGBCL-DH-*BCL2* ROI. Conversely, DZ-associated transcripts were significantly ($p < 0.05$) enriched in ROI of $IGH^{UND}$ HGBCL-DH-*BCL2*.

**Supplementary Figure S3. AID expression in $IGH^{+}$ and $IGH^{UND}$ HGBCL-DH-*BCL2***

(A) AID expression in representative $IGH^{+}$ and $IGH^{UND}$ HGBCL-DH-*BCL2*(-*BCL6*) cases, as measured by IHC. Histograms summarize frequencies of AID-expressing and-negative cases among $IGH^{+}$ and $IGH^{UND}$ subsets.

**Supplementary Figure S4. Protein levels of CD79B, and formation of complexes with IGH and CD79 in reactive lymphoid tissues and HGBCL-DH-*BCL2***

A) CD79B IHC analysis in representative reactive lymphoid tissues using the indicated monoclonal antibodies. Follicular B cells residing in the mantle zone show intense plasma membrane-associated CD79B immunoreactivity, in contrast to GCB cells expressing lower, mostly intracellular, CD79B levels.

B) Flow cytometric analysis of sCD79B in tonsillar CD19-gated naïve ($IGD^{+}CD38^{-}$), total GC ($IGD^{-}CD38^{+}$), GC-LZ ($IGD^{-}CD38^{+}CXCR4^{lo}$) and GC-DZ ($IGD^{-}CD38^{+}CXCR4^{hi}$) B cells. Histograms indicate CD79B median fluorescence intensity (MFI) values in each B cell subset.

C) IGG/CD79B protein complexes in a representative reactive GC, analyzed by PLA. GC DZ, LZ and mantle zone (M) areas from a rLN are labeled. Inset highlights PLA signals in GC LZ-restricted B cells. Quantification of $PLA^{+}$ cells is shown in Figure 4G.

D) IGM/CD79A protein complexes in two representative $IGM^{+}$ HGBCL-DH-*BCL2* measured by PLA. Insets highlight PLA signals within individual cells of representative tumor areas. Histogram depicts mean frequency (+/- SEM) of $PLA^{+}$ HGBCL-DH-*BCL2* (-*BCL6*) lymphoma B cells within n=5 fields of view.

E) CD79A/CD79B complexes in representative $IGH^{+}$ and $IGH^{UND}$ HGBCL-DH-*BCL2* cases, measured by PLA. Insets highlight PLA signals within individual cells of representative tumor areas.

F) Quantification of CD79A/B $PLA^{+}$ malignant B cells within n=5 fields of view for two $IGH^{+}$ and two $IGH^{UND}$ HGBCL-DH-*BCL2* (-*BCL6*) cases. Histogram depicts mean frequency (+/- SEM) of $PLA^{+}$ HGBCL-DH-*BCL2* (-*BCL6*) lymphoma B cells within n=5 fields of view (black, green and orange circles) in 4 cases.

(G) Reconstruction of the evolutionary trajectory of three $IGH^{UND}$ HGBCL-DH-*BCL2(-BCL6)* based on the comparison of clonal *IGHV* rearrangements with the preceding FL. Boxes indicate single nucleotide substitutions leading to amino acid replacements. Additional, conservative *IGHV* single nucleotide substitutions shared between HGBCL and FL, or unique to either of the metachronous tumors, were omitted for clarity.

**Supplementary Figure S5 Interpretation of *IGH* BAP and *IGH::BCL2* FISH signals**

(A) Representative patterns of *IGH* break apart (BAP) FISH signals in lymphomas with different configurations of *IGH* loci. Red and green bars indicate probes annealing to regions of *Chr14* mapping respectively 5’ and 3’ of the *IGH* locus.

(B) Representative *IGH::BCL2* dual fusion FISH signal patterns in HGBCL-DH-*BCL2*(-*BCL6*). Red and green bars indicate probes annealing respectively to *BCL2* and *IGH* loci. The lymphoma represented in the upper and lower right panels refers to a case where *MYC* and *BCL2* targeted in trans two *IGH* loci.

**Supplementary Figure S6. *RAG1/2* expression in IGH$^{UND}$ HGBCL-DH-*BCL2* cell line models**

(A) Predicted FISH signal patterns using *BCL2/BCL6* dual-fusion probes in healthy B cells and lymphomas carrying *BCL2* and *BCL6* rearrangements, targeting either the same genomic region (*cis*), or different ones (*trans)*. Colored bars refer to FISH probes.

(B) Representative FISH images with *BCL2/BCL6* dual-fusion probes, on metachronous FL (# 353) and HGBCL-DH-*BCL2-BCL6* (#245) cases described in Figure 6. The merge image (most right panels) indicates a pattern of probe hybridization suggestive of *BCL6* and *BCL2* rearrangements targeting different chromosomes.

(C) Normalized *RAG1/2* expression values in HGBCL-DH-*BCL2-BCL6* (#245) as compared to reactive lymph node (rLN), measured by RNA-seq.

(D) Normalized *RAG1/2* transcript levels in HGBCL-DH-*BCL2*(-*BCL6*) cell lines relative to 297 ALL cells, quantified by RT-PCR or RNA-sequencing ($^{\#}$). 293T and SU-DHL-5 cells were included as negative controls. Bars indicate mean values from n=3 biological replicates (+SEM).

(E) Representative (n $\geq$ 3) FACS analyses of sIGK/IGL expression in HGBCL-DH-*BCL2*, DLBCL and BL cell lines expressing *RAG1/2*. Lymphoma lines bearing t(8;22)(q24;q11) are indicated.

**Supplementary Figure S7. Inducible IgH extinction in λ-MYC mouse B cell lymphomas triggers RAG1/2-dependent BCR revision *in vivo***

(A) Interphase DNA FISH using break apart probes to capture *IGH*, *MYC*, *BCL2* and *BCL6* locus rearrangements in COH-DHL1 and COH-THL1 cell lines. Dual fusion DNA FISH was performed to capture *IGH*::*BCL2*, *IGH*::*MYC* and *cis/trans BCL6*::*BCL2* fusions in COH-DHL1/THL1 cell lines.

(B) Predicted structure of chromosomal rearrangements targeting *IGH, MYC, BCL2* and *BCL6* loci in COH-THL1 cells, as suggested by karyotype and FISH data.

(C) Class-specific IGH expression in representative COH-DHL1 and COH-THL1 cells from FFPE cytoblocks, measured by IHC.

(D) *IGKC* and *IGLC* transcripts in representative COH-DHL1 and COH-THL1 cells from FFPE cytoblocks, measured by RNA scope.

(E-F) Cre recombinase-induced BCR extinction in murine primary λ-MYC;B1-8f lymphoma cell lines according to[5]. Three weeks after intravenous inoculation of BCR-less lymphoma cells into immunocompetent syngeneic mice, animals were sacrificed and splenic cell suspensions assessed by FACS for MYC-expressing $BCR^+$/$BCR^-$ malignant B cells.

(F) Representative FACS analyses of pre-gated $CD19^+FSC^{hi}$ splenic lymphoma B cells from two lymphoma-transplanted animals inoculated three weeks earlier with $BCR^-$ tumoral cells. Recipient animal #1 showed expansion of $IgM^-$ (*B1-8Δ*) $MYC^+$ tumor cells. In recipient #2, a sizeable subset of $IgM^+$ $MYC^+$ tumoral cells co-existed with BCR-less counterparts.

(G) *Rag1/2* expression in ex vivo λ-MYC $BCR^+$ and $BCR^-$ lymphoma cells isolated from the bone marrow of recipient animals transplanted with either one of two independent tumor lines, measured by RNA-seq. (*** $p < 0.001$ t-test). Each circle represents a biological replicate.

(H) Cartoon depicting *Igh* loci captured in λ-MYC; B1-8f lymphoma B cells before and after Cre-mediated recombination, including tumor cells shifting from a $sBCR^-$ to a $sBCR^+$ phenotype (BCR-

>+). Arrows indicate PCR primers used to distinguish by genomic PCR *B1-8f* from wild-type *Igh* alleles.

(I) *Igh* genotyping of FACS-sorted IgM$^+$ MYC$^+$ malignant B cells retrieved from the spleen of recipient #2 (BCR- > +). The failure to amplify the *B1-8f* allele confirmed origin of the tumor cells from Cre-recombined BCR-less *B1-8* knock-out (Δ) lymphoma B cells. λ-MYC; B1-8f tumor cells prior to Cre-mediated recombination were included as positive control for the *B1-8f* allele (blue arrow).

(J) Examples of de novo productive *Ighv* rearrangements amplified by genomic PCR from BCR- > + (*B1-8*$^{\Delta}$) malignant B cells of recipient #2.
